# Carbon oxidation with sacrificial anodes to inhibit O_2_ evolution in membrane-less bioelectrochemical systems for microbial electrosynthesis

**DOI:** 10.1101/2022.08.23.504965

**Authors:** Nils Rohbohm, Tianran Sun, Ramiro Blasco-Gómez, James M. Byrne, Andreas Kappler, Largus T. Angenent

## Abstract

Microbial electrosynthesis is an emerging biosynthesis technology that produces value-added chemicals and fuels and, at the same time, reduces the environmental carbon footprint. However, constraints, such as low current densities and high inner resistance, disfavor this technology for industrial-scale purposes. The cathode performance has been strongly improved in recent years, while the anode performance has not been given enough attention despite its importance in closing the electric circuit. For traditional water electrolysis, O_2_ is produced at the anode, which is toxic to the anaerobic autotrophs that engage in microbial electrosynthesis. To overcome O_2_ toxicity in conventional microbial electrosynthesis, the anode and the cathode chamber have been separated by an ion-exchange membrane to avoid contact between the microbes and O_2_. However, ion-exchange membranes increase the maintenance costs and compromise the production efficiency by introducing an additional internal resistance. Furthermore, O_2_ is inevitably transferred to the catholyte due to diffusion and electro-osmotic fluxes that occur within the membrane. Here, we proved the concept of integrating carbon oxidation with sacrificial anodes and microbes to simultaneously inhibit the O_2_ evolution reaction (OER) and circumvent membrane application, which allows microbial electrosynthesis to proceed in a single chamber. The carbon-based anodes performed carbon oxidation as the alternative reaction to the OER. This enables microbial electrosynthesis to be performed with cell voltages as low as 1.8-2.1 V at 10 A·m^-2^. We utilized *Methanothermobacter thermautotrophicus* ΔH in a single-chamber Bioelectrochemical system (BES) with the best performing carbon-based anode (*i*.*e*., activated-carbon anode with soluble iron) to achieve a maximum cathode-geometric CH_4_ production rate of 27.3 L·m^-2^·d^-1^, which is equal to a volumetric methane production rate of 0.11 L·L^-1^·d^-1^ in our BES, at a coulombic efficiency of 99.4%. In this study, *Methanothermobacter thermautotrophicus* ΔH was majorly limited by sulfur that inhibited electromethanogenesis. However, this proof-of-concept study allows microbial electrosynthesis to be performed more energy-efficiently and can be immediately utilized for research purposes in microbial electrosynthesis.

## INTRODUCTION

Globally increasing demands for value-added chemicals and fuels require innovative chemical and biochemical synthesis strategies to proceed in a more cost-efficient and environmentally friendly manner. Microbial electrosynthesis is a bioprocess with bioelectrochemical systems to convert inorganic gases into valuable, multi-carbon compounds to store excessive renewable energy such as wind or solar energy. ^1, 2^ This makes it an attractive technology for achieving the goal of a carbon-neutral society. Microbial activity and electrode performance are critical factors in determining the overall efficiency of microbial electrosynthesis. However, the inherent problem of scaling-up microbial electrosynthesis is the currently low current density of the bioelectrochemical system, which results in a considerably lower production rate compared to chemical electrosynthesis that produces the same compounds (**Table S1**). To increase the current density, the electrochemical performance must be increased by, for example, improving the cathode performance. Indeed, several studies have focused on the modification of the cathode to improve microbe-electrode interactions and its biocompatibility.^3, 4^

Besides the biocompatibility problems, another troublesome part of bioelectrochemical systems is the ion-exchange membrane. Classically, the anode is separated from the cathode by an ion-exchange membrane to avoid interaction of the O_2_ evolving at the anode with the strict anaerobic microbes in the cathodic chamber.^5^ However, applying an ion-exchange membrane in bioelectrochemical systems has several disadvantages. By including an ion-exchange membrane, the internal resistance is increased.^6^ This will lead to decreased ion transport, which consequently limits electron transfer and reduces the current generation. Ideally, protons are primarily transported from the anodic to the cathodic chamber to maintain hydrogen production. However, the transfer of other cations, such as sodium or potassium, to the cathode chamber cannot be neglected and reduces the ionic transport number of protons. Consequently, the slower mass transfer rate of protons leads to a pH gradient.^7^ In addition, the ion-exchange membrane also generates an electro-osmotic flow that allows all ionic and non-ionic solutes, including O_2_, to pass through the cathode chamber, thus, making its purpose ineffective.^8^ The pore size of the membrane in addition to O_2_ permeation, which allows biofilm formation, makes it vulnerable to fouling.^9^ Finally, the capital costs for membranes are high, and combining all the other disadvantages will result in high maintenance and operating costs.^10, 11^

The ideal bioelectrochemical system would, therefore, be a single-chamber system in which the ion-exchange membrane has been removed.^12^ This would improve not only the current generation but also the synthesis efficiency. Cl_2_ and O_2_ are the main gases produced on the anode side, with Cl_2_ being not only a strong oxidizing agent to many elements but also very toxic. While Cl_2_ can be avoided by removing the chlorine-ions from the anodic chamber, O_2_ would still be produced without modifications at the anode through water splitting and hinder anaerobic microbes from growing for microbial electrosynthesis to proceed. Thus far, few reports have been published showing membrane-less bioelectrochemical systems, and most reports are about microbial fuel cells and enzymatic fuel cells. Furthermore, most membrane-less, single-chamber systems were operated without O_2_-sensitive microbes.^13, 14^ Giddings et al.^5^ designed a membrane-less system for microbial electrosynthesis in which the anode was placed at the boundary of the liquid interface to avoid O_2_ from mixing. Besides, different patent applications for single-chamber bioelectrochemical systems were filed with setups to capture and remove O_2_ away from the cathode.^15, 16^ However, none of these innovations had tried to circumvent the O_2_-evolution reaction (OER) at the anode.

An appealing approach to avoid O_2_ formation at the anode is carbon-assisted water electrolysis (CAWE) for which carbon is oxidized at the anode instead of the OER in conventional electrolysis.^17, 18^ The classical hydrogen-evolution reaction (HER) is not disturbed and proceeds at CAWE. CAWE can include: **(1)** the oxidation of carbon-based electrodes, which we refer to as carbon oxidation with sacrificial anodes; and **(2)** the oxidation of soluble carbon materials (which we did not study) or non-soluble carbon materials in the anolyte. Here, we address carbon oxidation of sacrificial anodes, forming CO_2_, CO, and oxygenated surface-functional groups. The desired two chemical reactions can be described as follows (**Eq. 1** and **Eq. 2**):

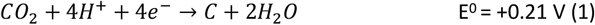

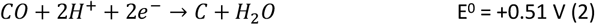

A major advantage of using CAWE instead of conventional water electrolysis with O_2_ evolution at the anode is the lower operating cell voltage, resulting in lower energy consumption rates. For example, the standard electrode cell potential for the four-electron process (**Eq. 1**) for CAWE is E^0^_cell,CAWE_ = -0.21 V, while the conventional water electrolysis reaction requires E^0^_cell,water electrolysis_ = -1.23 V.^17^ Another advantage of CAWE compared to conventional water electrolysis is the higher carbon-oxidation reaction rate than the OER rate at the anode (exchange current density: j_0,CAWE_ =10^−6^ A·cm^-2^ *vs*. j_0,OER_ = 10^−7^ A.cm^-2^).^17^ Albeit, the carbon-oxidation reaction at the anode is still slower than the HER rate at the cathode (j_0,HER_ = 10^−3^ A.cm^-2^).^19^ Finally, a necessary substrate for microbial electrosynthesis is CO_2_ or CO, which is the desired product of carbon oxidation with sacrificial anodes. Thus, the matching product formation as extra carbon source for the microbes is another pertinent advantage of carbon oxidation with sacrificial anodes for microbial electrosynthesis.

An example of carbon oxidation with carbon-based anodes was shown by Holubowitch et al.^18^, who demonstrated an abiotic membrane-less electrochemical cell that was capable of reducing O_2_ to deoxygenate water. While their main focus was on reducing O_2_, we identified with their data that either no O_2_ was produced or O_2_ production was decreased at the anode and that O_2_ was electrochemically reduced at the cathode. Carbon oxidation with carbon-based anodes was not only tested abiotically but also for bioelectrochemical systems. Lai et al. ^20^ found a lower O_2_ content and a higher CO_2_ content in their anodic compartment using graphite. They concluded that carbon oxidation with the graphite anode was the main cause of the lower O_2_ formation. A follow-up study for the removal of perchloroethylene in a membrane-less bioelectrochemical system used graphite granules as a single anode to avoid O_2_ production.^21^ To our knowledge, carbon oxidation with sacrificial anodes has not been tested for microbial electrosynthesis.

The objective of this work was a proof of concept to show that carbon oxidation can be utilized in a single-chamber bioelectrochemical system to specifically perform microbial electrosynthesis. We used different types of carbon electrodes as anodes and assessed whether O_2_ formation was inhibited or just decreased. In addition, we tested iron (soluble and coated) materials with carbon electrodes to further circumvent the OER by including a parallel Faradaic reaction. After we had studied the feasibility of our carbon electrodes and deciphered the mechanism of the inhibition, we operated our carbon electrodes in bioelectrochemical systems to evaluate their potential for microbial electrosynthesis. Here, we report the successful implementation of carbon oxidation with a sacrificial anode for microbial electrosynthesis using *Methanothermobacter thermautotrophicus* ΔH, which is an autotrophic and anaerobic microbe, and which is being used in the power-to-gas concept to store renewable electric power into CH_4_ gas.^22^

## RESULTS AND DISCUSSION

### The minimal required cell potential is lower for carbon oxidation than for O_2_ evolution at the anode, while the kinetics of the carbon oxidation reaction is higher

For our proof-of-concept process, we used the two carbon-oxidation reactions from CAWE compared to the OER from conventional water electrolysis. The two carbon-oxidation reactions have a theoretically more favorable thermodynamic potential of +0.21 V *vs*. RHE (reversible hydrogen electrode which depends on the solution pH) for CO_2_ production (four electrons in **Eq. 1**) and +0.51 V *vs*. RHE for CO production (two electrons in **Eq. 2)**, respectively^23^, than the OER, with a standard potential of +1.23 V *vs*. RHE. For conventional water electrolysis with the OER, the overall cell potential would be -1.23 V. However, electrode materials have properties that introduce an overpotential. In the case of carbon electrodes, the overpotential can be up to 500 mV or higher.^24^

To measure the overpotentials (η) for our specific BES for carbon oxidation with carbon-based anodes, OER, and HER, we performed linear sweep voltammetry on the activated carbon (as the anode) and carbon cloth (as the cathode). Linear sweep voltammograms demonstrated that the measured overpotential (│η│ = [E_onset potential_ + E_reference electrode correction_] – [E^0^ - 0.059·pH]) of carbon oxidation with activated carbon was +1.16 V, while the HER for carbon cloth had an overpotential of +0.190 V (**Fig. 1**). Thus, the minimum required cell potential for CAWE anodes would be -1.56 V (E^0^_HER_ - η_HER_ - [E^0^_carbon oxdiation_ + η_carbon oxidation_]), which is -1.37 V greater than the theoretical value (E^0^_cell, carbon oxidation_ = E^0^_HER_ - E^0^_carbon oxidation_ = -0.21 V). The OER with the activated carbon had an overpotential of +0.86 V (**Fig. 1**). Therefore, the minimal required cell potential for conventional water electrolysis would be equivalent to -2.28 V (E_cell, water electrolysis_ = E^0^_HER_ - η_HER_ - [E^0^_OER_ + η_OER_]), which is -1.05 V greater than the theoretical value. The considerably less negative overall cell potential for CAWE compared to conventional water electrolysis should lower the energy consumption in our system as well.

**Figure 1:**
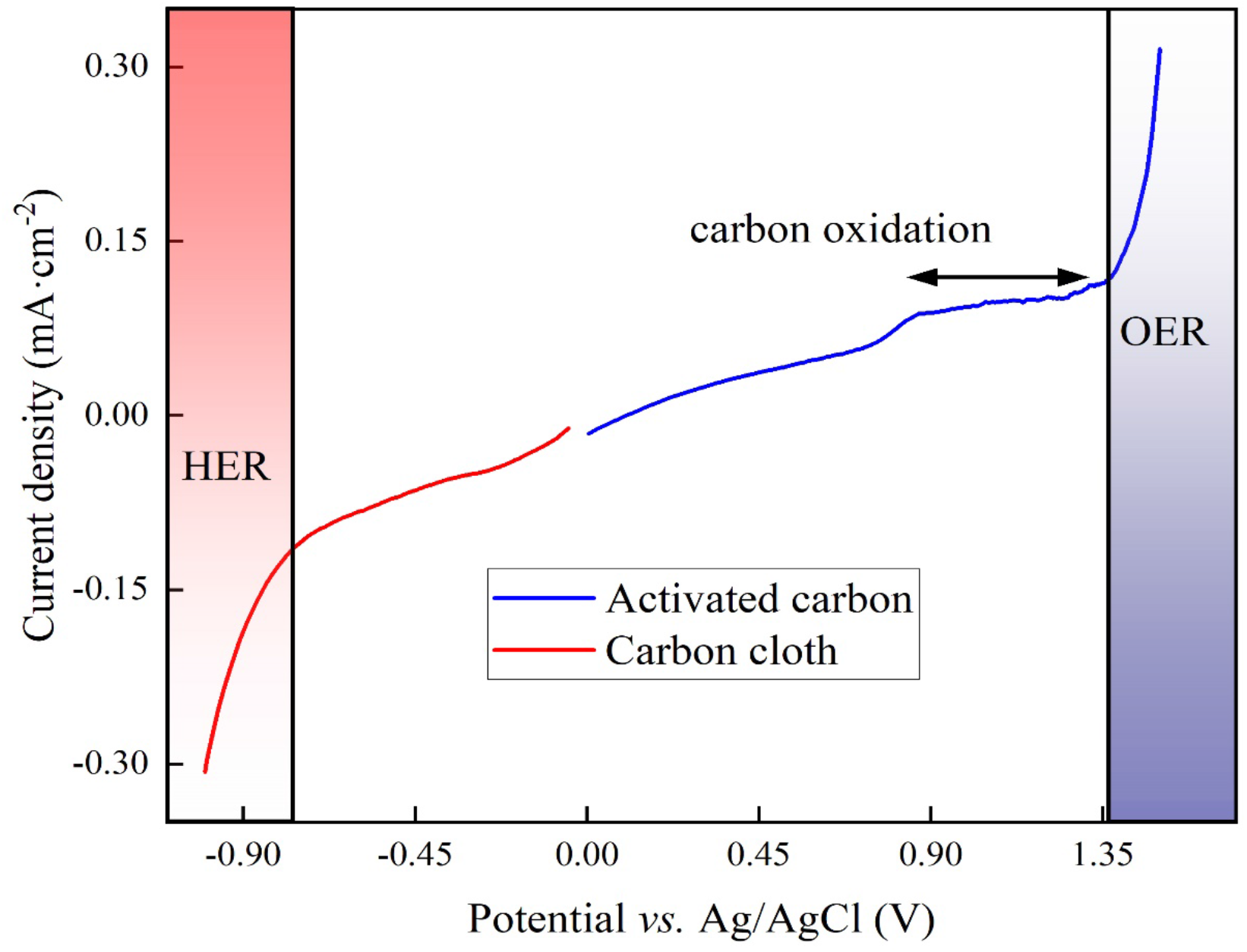
Linear sweep voltammogram of the activated-carbon anode (blue line) and the carbon-cloth cathode (red line) in 66 mM phosphate buffer (pH = 7.2) at a scan rate of 20 mV·s^-1^.

However, besides the cell potentials, the kinetics of the different electrochemical reactions are also important. The linear-sweep voltammogram of activated carbon did not show any pristine carbon-oxidation peaks to compare the kinetic value of the carbon oxidation to the OER (**Fig. 1**). However, the exchange current density, which is a dimension of the reaction rate, can be deduced from the linear-sweep voltammograms by means of the Butler-Volmer equation (**Eq. S1** in supporting note 1 of the SI).^25^ The exchange current density for carbon oxidation was 10^−5^ A·cm^-2^ and was one magnitude higher than the OER at 10^−6^ A·cm^-2^. Therefore, the carbon oxidation reaction with sacrificial anodes is a faster electrochemical reaction than OER.

### Carbon-based anodes with and without iron species circumvented the need for a membrane, opening the possibility of a membrane-less system

To identify whether O_2_ production is inhibited by carbon oxidation, we measured the O_2_ content of an abiotic, two-electrode electrochemical system for five days. The O_2_ content measurement was performed abiotically to circumvent using L-cysteine, which would be the preferred sulfur source and reducing agent in our medium for the BES experiments, and which would have removed O_2_ from the electrolyte. The abiotic experiments were performed in triplicate for two temperature conditions: room temperature and 60°C, which is the optimal growth temperature of *Methanothermobacter thermautotrophicus* ΔH. We used a graphite rod, thermally activated graphite felt, carbon cloth, and activated carbon as our carbon anodes for four out of seven treatments and included a platinum anode that was poised at 10 A·m^-2^ in 0.066 M phosphate buffer as the positive control to compare the results of the carbon oxidation with sacrificial anodes to the OER (**Fig. 2**).

**Figure 2:**
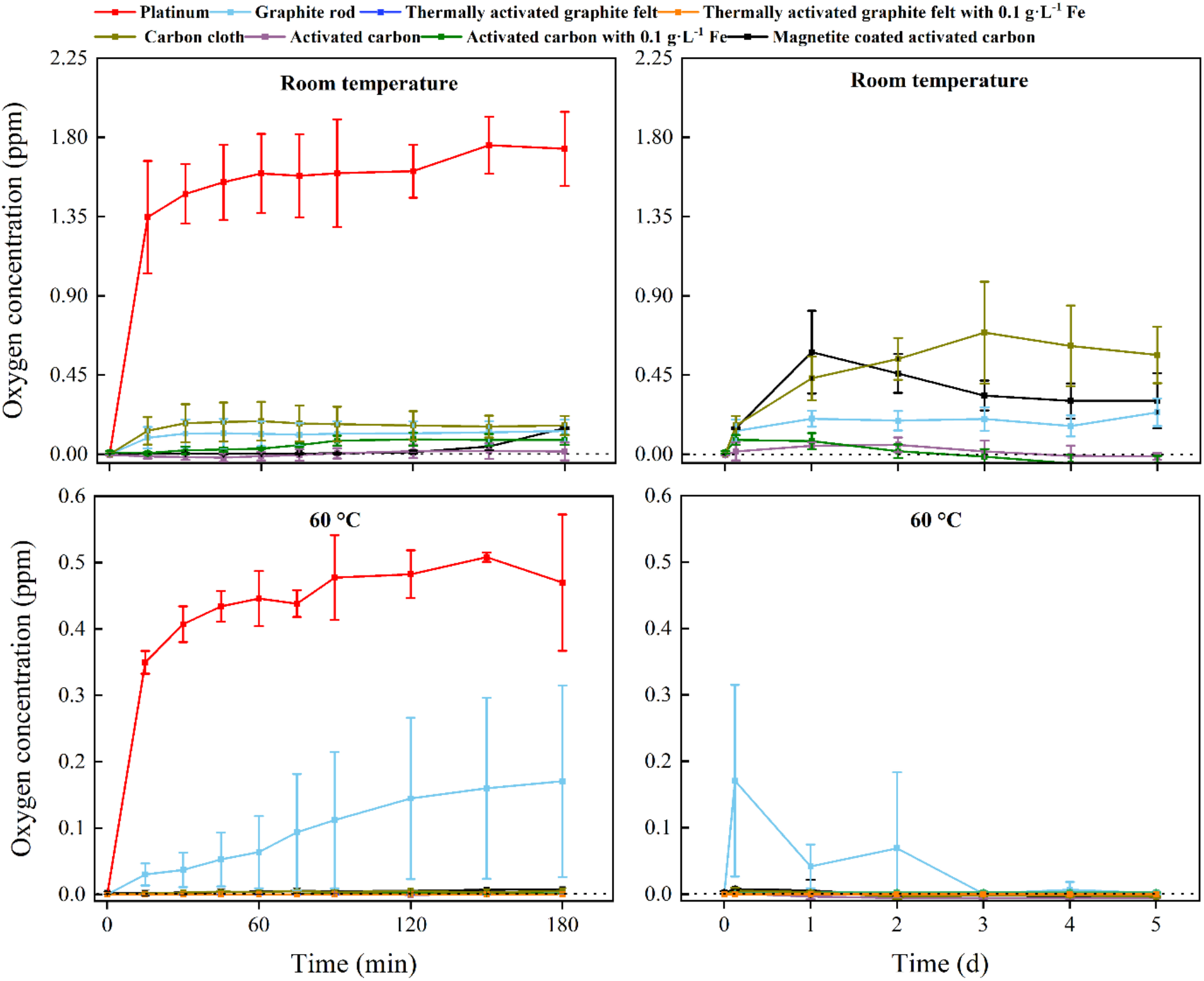
O_2_ concentration in the electrolyte of platinum, graphite-rod, thermally activated graphite felt, carbon-cloth, and activated-carbon anodes over time. Two temperature conditions were included. Two treatments with thermally activated graphite felt and three treatments with activated carbon were included, including iron additions. The error bars show the standard deviation for the average date from triplicate systems: **(A)** the O_2_ concentration at room temperature for 180 min; **(B)** the O_2_ concentration at room temperature for 5 days; **(C)** the O_2_ concentration at 60°C for 180 min; and **(D)** the O_2_ concentration at 60°C for 5 days. We omitted the data for the first 180 min from panels B and D for clarity reasons. No L-cysteine was added to these abiotic experiments.

We added Fe^2+^ species in the form of: **(1)** soluble Fe^2+^ species (0.1 g.L^-1^ of Fe^2+^, corresponding to 0.27 g·L^-1^ FeSO_4_); and **(2)** coated magnetite for three out of seven treatments to this abiotic O_2_ evolution study. We added Fe^2+^ species because abiotic electrochemical systems increased the current density for carbon oxidation with sacrificial anodes without reducing the Coulombic efficiency.^26^ Soluble Fe^2+^ is directly reduced at the carbon-based anode to Fe^3+^ at electrochemical rates that are higher than carbon oxidation and the OER (**Eq. 3**).

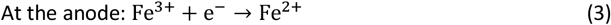

Therefore, soluble Fe^2+^ is an ultimate electron donor in addition to the carbon-based anodes when Fe^2+^ is continuously added to the system. Soluble Fe^2+^ was added to the thermally activated-graphite-felt and activated-carbon anodes. Magnetite is another biocompatible Fe^2+^ source. We coated magnetite on the activated carbon to form the anode after performing a magnetite-to-carbon ratio analysis (supporting information note 2 of the SI). Magnetite is a mixed-valent Fe(II), Fe(III) magnetic iron mineral that oxidizes to maghemite (γ-Fe_2_O_3_), donating one electron (**Eq. 4**):

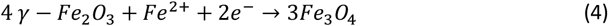

All carbon-based anodes with and without Fe^2+^ species showed an inhibitory effect on the O_2_ production compared to the positive control for the first 180 min at room temperature, with the graphite-rod and carbon-cloth anodes producing more O_2_ compared to the thermally activated-graphite-felt and activated-carbon anodes (**Fig. 2A**). Thus, a pretreatment to activate the graphite or carbon will permit a faster inhibition of O_2._ We turned off the electrochemical system with the platinum anode after 180 min to avoid accumulating O_2_ and H_2_ and creating a *Knallgas* reaction. For the next 5 days, the accumulated O_2_ concentration of all the carbon-based anodes was relatively low (<1 ppm), but the thermally activated-graphite-felt and activated-carbon anodes with and without soluble Fe^2+^ could sustain a nearly O_2_-free environment (**Fig. 2B**). However, magnetite-coated activated-carbon anodes showed a higher O_2_ concentration at room temperature compared to the other activated carbon-based electrodes for unknown reasons (**Fig. 2B**).

In contrast, all the experiments with carbon-based anodes at 60°C inhibited O_2_ production during the first 180 min, except for the graphite-rod anode (sky-blue line, **Fig. 2C**). In addition, no O_2_ was produced at 60°C for the thermally activated-graphite-felt and activated-carbon anodes with and without soluble Fe^2+^ and activated-carbon anodes with magnetite (**Fig. 2D**). Because the kinetics of a reaction are directly related to the temperature, the increase in the temperature, therefore, increased the carbon-oxidation rate. Likewise, the kinetics of the OER were also affected by the increase in temperature. However, our results show that no production of O_2_ occurred and that carbon oxidation was the dominating reaction in our system (**Fig. 2D**).

During the five-day operating period, the O_2_ concentration in the electrolyte with the graphite-rod anode gradually decreased, suggesting that graphite needs an activation time to oxidize its surface and efficiently perform carbon oxidation (**Fig. 2D**). Indeed, the thermally activated-graphite-felt and activated-carbon anodes did not show oxygen evolution during the first couple of days of the operating period (**Fig. 2D**). The concentrations of CO_2_ and CO were measured for the two temperature conditions and the seven treatments, but the concentrations were very low for both gases due to the low applied current of up to 1 mA cm^-2^ and were not further monitored. In summary, the carbon-based anodes circumvented O_2_ production at 60°C and were used for further thermophilic BES experiments. Because we completely inhibited O_2_ evolution at the anode, we can remove the membranes from the bioelectrochemical system to perform microbial electrosynthesis. When comparing these abiotic results to the BES experiments, the abiotic O_2_ evolution results will be conservative because we added reducing agents only to the microbial medium.

### Different carbon materials as the anode affected the CH_4_ production duration of microbial electrosynthesis, and competitive methane production rates were achieved to other BES studies

In our proof-of-concept BES for microbial electrosynthesis, we opted for the microbe *M. thermautotrophicus* ΔH due to its high growth rate (doubling time of 120 min) and relatively high CH_4_ production rate.^22^ This archaeal strain grows anoxically at 60°C and shows a resilient behavior in the presence of O_2_ by having the ability to grow after exposure to air for several hours.^27^ However, as a strict anaerobic methanogen, it is still sensitive to O_2_ for longer operating periods, resulting in the absence of growth when prolonged O_2_ is present. The BES experiments were performed at 60°C and close to neutral-pH conditions with graphite-rod, thermally activated-graphite-felt, and activated-carbon anodes in an experimental design with six treatments including soluble Fe^2+^ and magnetite. We set the current at 2 mA and with a 1x1 cathode-surface area the cathode-based geometric current density was 1 mA cm^-2^ (**Fig. 3**).

**Figure 3:**
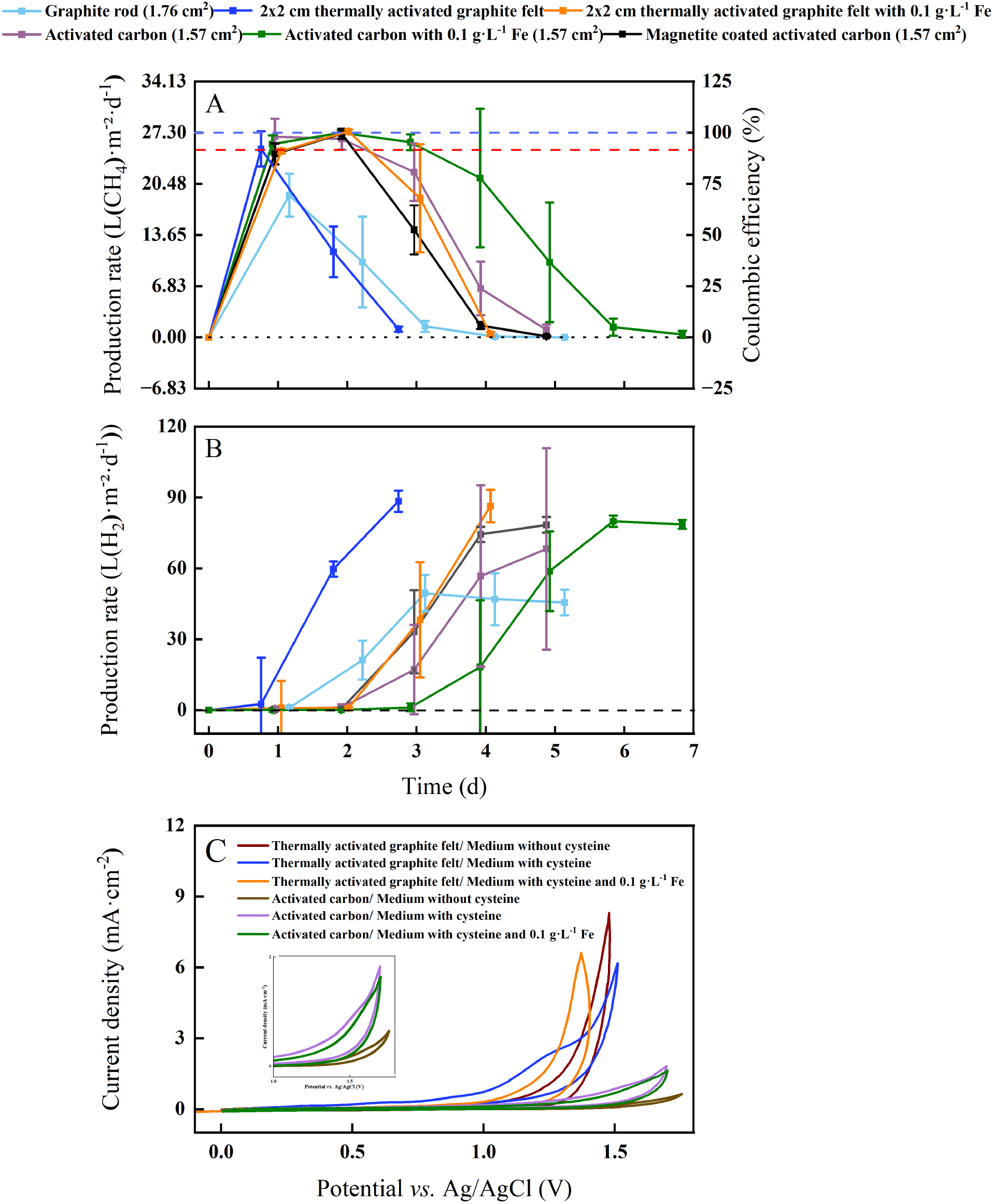
Membrane-less BES performance for electromethanogenesis and cyclic voltammetry analysis with L-cysteine: **(A)** CH_4_ production rates corrected to the cathode surface area and Coulombic efficiencies (at I = 10 A·m^-2^); **(B)** H_2_ production rate corrected to the cathode surface area during batch experiments over time with the thermophilic archaeon M. thermautotrophicus ΔH. Graphite-rod, thermally activated graphite felt and activated-carbon anodes were included with two treatments and three treatments for thermally activated graphite felt and activated carbon, respectively, including iron additions (soluble iron and magnetite). The error bars show the standard deviation for the average data from triplicate systems. For **(A)** and **(B)** Carbon cloth was used as the cathode: (Y-axis 1) cathode-based geometric CH_4_ production rates and H_2_ production rates for six treatments; and (Y-axis 2) for **(A)** Coulombic efficiencies for six treatments. After reaching a maximum H_2_ production rate, we stopped the current to not accumulate excessive amounts of H_2_ in our BESs. While the blue-dotted line represents 100% Coulombic efficiency at a 27.30 L.m^-2^·d^-1^ methane production rate, the red-dotted line shows the empirical cathode-based geometric CH_4_ production rate of > 25 L.m^-2^·d^-1^ to determine the duration of the optimum performance period; and **(C)** cyclic voltammograms of thermally activated graphite felt and activated carbon at a scan rate of 5 mVs^-1^ in MS-medium. In total 6 cyclic voltammogramms were recorded where the MS-medium for each carbon-based electrode contained L-cysteine without soluble Fe^2+^; L-cysteine with soluble Fe^2+^; and without L-cysteine and without soluble Fe^2+^

*M. thermautotrophicus* ΔH produced CH_4_ with CO_2_ as the sole carbon source during carbon oxidation with sacrificial anodes for all treatments (**Fig. 3A**). After an operating period of one day, the maximum cathode-based geometric CH_4_ production rates for the BESs with the graphite-rod, the thermally activated graphite felt, the activated-carbon anode were 18.97, 25.13, and 26.75 L·m^-2^·d^-1^, respectively (**Table 1**). This resulted in a maximum volumetric CH_4_ production rate of 0.1 L·L^-1^·d^-1^ for the BES with the activated-carbon anode. Coulombic efficiencies of 97.96% for the BES with the activated-carbon anodes suggest an efficient H_2_ uptake by the microbes at the cathode (**Fig. 3)**.

**Table 1.** Performance data of the carbon-based anodes.

| Carbon-based anode | Maximum geometric CH<sub>4</sub> production rate in L·m<sup>-2</sup>·d<sup>-1</sup> | Maximum volumetric CH<sub>4</sub> production rate in L·L<sup>-1</sup>·d<sup>-1</sup> | Optimum performance period in d |
| --- | --- | --- | --- |
| Graphite rod | 18.97 ± 2.84 | 0.07 ± 0.01 | 0 |
| Thermally activated graphite felt | 25.13 ± 2.34 | 0.10 ± 0.009 | 1 |
| Thermally activated graphite felt with 0.1 g·L<sup>-1</sup> of Fe<sup>2+</sup> | 27.48 ± 0.26 | 0.11 ± 0.001 | 2 |
| Activated carbon | 26.75 ± 2.38 | 0.10 ± 0.009 | 2 |
| Activated carbon with 0.1 g·L<sup>-1</sup> of Fe<sup>2+</sup> | 27.30 ± 0.29 | 0.11 ± 0.001 | 3 |
| Magnetite-coated activated carbon | 27.15 ± 0.65 | 0.11 ± 0.002 | 2 |

By adding soluble Fe^2+^ to thermally activate-graphite-felt and activated-carbon anodes at 60°C with *M. thermautotrophicus* ΔH, we achieved a higher maximum cathode-based geometric CH_4_ production rate of 27.48 and 27.30 L·m^-2^·d^-1^, respectively, resulting in a volumetric CH_4_ production rate of 0.11 L·L^-1^·d^-1^ for both and a higher Coulombic efficiency of 100.6% and 99.36% with the added soluble Fe^2+^ than without (**Table 1, Fig. 3A**). However, with magnetite-coated activated-carbon anode material, the methane production was inferior compared to activated-carbon anode material. The maximum cathode-based geometric CH_4_ production rate was 27.15 L·m^-2^·d^-1^m with a Coulombic efficiency of 99.40% (**Table 1** and **Fig. 3)**.

To compare the different experimental conditions in our study, we choose an empirical cathode-based geometric CH_4_ production rate of > 25 L·m^-2^·d^-1^ to determine the duration of the optimum-performance period for each treatment. Under this definition of the optimum performance period, the graphite-rod anode never reached this CH_4_ production rate, resulting in an optimum performance period of zero (**Table 1**). We achieved an optimum-performance period of 1 day for the thermally activated-graphite-felt anode. After an operating period of two days, the cathode-based geometric CH_4_ production rates for the BESs with the activated-carbon anode was 26.44 ± 1.40 L·m^-2^·d^-1^ after which it decreased, resulting in an optimum-performance period of two days (**Table 1**). By adding soluble Fe^2+^ to thermally activated-graphite-felt and activated-carbon anodes, the optimum-performance period was elongated to two and three days at an average cathode-based geometric CH_4_ production rate of 26.17 ± 0.29 and 26.36 ± 0.84 L·m^-2^·d^-1^, respectively (**Table 1**). However, with magnetite, the optimum performance period was shortened to two days at an average cathode-based geometric CH_4_ production rate of 25.81 ± 1.01 L·m^-2^·d^-1^, which is one day shorter than for the activated-carbon anodes with soluble Fe^2+^ (**Table 1**). For the activated-carbon anodes as an example, the pH remained between 7.3 and 7.8 (**Fig. S1**), and CO_2_ was never limited (**Fig. S2)**.

We achieved a maximum cathode-based geometric CH_4_ production rate of 27.48 L·m^-2^·d^-1^ with thermally activated graphite felt and 0.1 g L^-1^ soluble Fe^2+^, which was slightly higher than Kracke et al.^4^, who achieved a maximum of 26.76 L·m^-2^·d^-1^ at a cathode-based geometric current density of 1 mA cm^-2^. The authors utilized a mesophilic methanogen (*i*.*e*., *Methanococcus maripaludis*) in a BES with membranes and a NiMo cathode. However, the reported maximum volumetric CH_4_ production rate by Kracke et al.^4^ was 1.38 L·L^-1^·d^-1^, which was 12.5 times higher than the 0.11 L·L^-1^·d^-1^ for our study due to a more optimum electrode-surface-to-reactor-volume ratio. An even higher volumetric CH_4_ production rate of 2.9 ± 1.2 L·L^-1^·d^-1^ compared to Kracke et al. ^4^ was achieved by Baek et al.^28^, using a zero-gap electrochemical cell. Despite its lower geometric CH_4_ production rate of 18.25 L·m^-2^·d^-1^ and side-products of acetate and propionate (at a cathode-based geometric current density of 1.74 mA cm^-2^)^28^, the zero-gap electrochemical cell has an optimized electrode-surface-to-reactor-volume ratio and has the propensity to increase the production rates for microbial electrosynthesis, including for our system.

Noteworthy is the considerably lower volumetric CH_4_ production rates for all BES studies compared to gas fermentation by sparging H_2_ and CO_2,_ for which volumetric CH_4_ production rates of 163 L·L^-1^·d^-^ _1_ had already been achieved in the 1990s._29_ It is clear that several breakthroughs are necessary to close the gap between microbial electrosynthesis and gas fermentation for CH_4_ production. Regardless, our proof-of-concept was successful for which we could produce the highest published cathode-based geometric CH_4_ production rate of 27.48 L·m^-2^·d^-1^ with our membrane-less BES with sacrificial anodes.

### The duration of methane production in the single-chamber bioelectrochemical system was only a few days due to sulfur oxidation at the anode and a resulting sulfur limitation

Our abiotic O_2_ measurements with graphite-rod anodes showed O_2_ production at 60°C (**Fig. 2D**), identifying that O_2_ inhibition to the thermophilic methanogen was a reason for a zero-day optimum-performance period during our BES experiment with graphite-rod anodes (**Fig. 3A**). The thermally activated-graphite-felt anodes and activated-carbon anodes during abiotic experiments exhibited no O_2_ production after an operating period of five days (**Fig. 2D**), explaining the longer optimum-performance period for these anodes than the graphite-rod anodes (**Fig. 3A**). Still, we could only maintain a longest optimum-performance period of three days for the BES with activated-carbon anodes and soluble Fe^2+^ (**Table 1**). By comparing the CH_4_ production rate (**Fig. 3A**) with the H_2_ production rate (**Fig. 3B**), we observed different onsets of H_2_ production after the CH_4_ production had peaked for graphite-rod, thermally activated-graphite-felt, and activated-carbon anodes (with and without iron). This indicated that the electrochemical system was not limiting (H_2_ production did not stop). Thus, carbon oxidation had not diminished the available carbon at the anode to maintain a working electrochemical system.

This also became clear when we increased the size of the anode surface areas and observed a similar optimum-performance period of one day for each of the BESs with the 1x1 cm-, 1x2 cm-, and 2x2 cm-thermally activated-graphite-felt anodes and with L-cysteine as the sulfur source and reducing agent in our growth medium (**Fig. S3**). While the larger electrode provided a larger surface area for the carbon oxidation to occur and a more stable cell voltage (**Fig. S4**), it did not improve the performance. Such a similar performance showed that, indeed, the carbon material was not limiting but that something else was limiting the methanogens to produce CH_4_. Without a membrane, L-cysteine oxidizes to L-cystine^30^, which may render L-cysteine unavailable as a sulfur source and reducing agent. Therefore, we investigated whether the relatively fast oxidation of reducing sulfur components in the growth medium would diminish our sulfur source for methanogenic growth and activity.

Instead of L-cysteine, we added 0.1 g·L^-1^ Na_2_S as a sulfur source and reducing agent to our treatment for the BESs to have a measurable sulfur source and a more potent reducing agent to lower the oxygen redox potential (ORP) without harming the microbes.^31^ We used thermally activated-graphite-felt anodes with 0.1 g L^-1^ Fe^2+^ because this resulted in the maximum cathode-based geometric CH_4_ production rate (**Table 1**). However, we increased the anode size from 1x1 to 2x2 cm and also increased the number of methanogenic cells during inoculation by 10 times compared to previous experiments. We compared the treatment with Na_2_S and soluble Fe^2+^ to the control with L-cysteine and soluble Fe^2+^, and obtained a similar: (1) CH_4_ production pattern; and (2) optimum-performance period of two days compared to previous results (**Fig. 3A**), and between the treatment and the control (**Fig. S5A**).

To determine whether sulfide was oxidized and limiting the methanogens in our BESs, the sulfide concentration of each sampling point was measured for the treatment BESs through a photometric assay. The sulfide concentration decreased rapidly after the first day, proving that sulfide was removed, likely by oxidation (**Fig. S5B**). After three days, sulfide was nearly depleted (**Fig. S5B**). In addition, the ORP value increased from -375 mV to -100 mV showing that the reducing power of sulfide was removed at the same time (**Fig. S5C**). These results suggest that the previous experiments were all limited in sulfur, such that the sulfur source and/or the reducing power became limiting to the methanogens (**Table 1**).

A potential inhibiting factor for *M. thermautotrophicus* ΔH is an ORP value that is not negative enough to provide reducing power. In fact, electromethanogenesis would already be inhibited at a minimum ORP of -200 mV.^32^ Because both sulfur compounds were oxidized during the BES operating conditions, the reducing power and, subsequently, the ORP value for methanogenesis should worsen. To find out whether the depletion of the sulfur source or the increase in the ORP due to the absence of the reducing agent limited the methanogens after sulfur oxidation, we added another treatment. Besides the Na_2_S and soluble Fe^2+^, we added Ti-NTA, L-cysteine, and soluble Fe^2+^. In fact, we introduced Ti-NTA at the beginning of the experiment and after every sampling point. However, no difference in the cathode-based geometric CH_4_ production rate was observed between Ti-NTA and the control (L-cysteine) (**Fig. S5A**). In addition, the ORP values were similar throughout the operating period and had crossed the minimum value of -200 mV between Day 1 and Day 3 (**Fig. S5C**). Thus, adding Ti-NTA did not lower the ORP because Ti^3+^ was also oxidized at the anode, preventing its reducing power from being effective. Therefore, we could not distinguish whether the depletion of the sulfur source or the increase in the ORP due to the absence of the reducing agent limited the methanogens.

Regardless, sulfur was the limiting factor of the single-chamber BES. Therefore, the recovery of sulfide *in situ* after sulfur oxidation is a desirable objective to circumvent the continuous supply of sulfide to the electrochemical system. For a recent study by Izadi et al.^33^, sulfide was oxidized at the anode to elemental sulfur after which the sulfide was recovered at the cathode by switching the polarity of the electrodes. Based on their study, we performed a BES experiment using thermally activated graphite felt as both anode and cathode while adding 0.1 g·L^-1^ Na_2_S as an additional sulfur source. The polarity of the electrodes was switched every 4 h to reduce the elemental sulfur to sulfide. However, our study did not have a desirable outcome because the cathode-based geometric CH_4_ production rate was lower than in all other BES experiments, possibly due to a stressed environment (**Fig. S6A**). In addition, the sulfide concentration was depleted faster, limiting CH_4_ production (**Fig. S6B**). We cannot rule out that sulfide was oxidized to higher charged sulfur oxides, such as sulfate or thiosulfate, which are more difficult to reduce back to sulfide, preventing the capability to recover sulfide.

### Soluble iron elongated the optimum-performance period for reasons that are related to sulfide oxidation

We performed cyclic voltammetry (CV) measurements with two carbon materials at the anode in an abiotic experiment with 6 treatments of which with and without L-cysteine and with and without soluble Fe^2+^ were included (**Fig. 3C**). We were interested in observing the onset potential of the sulfur oxidation during the forward electrochemical scan (5 mV s^-1^). The thermally activated graphite felt with L-cysteine and without soluble Fe^2+^, showed the earliest onset potential of the six cyclic voltammograms and an electrochemical shoulder with a mid-point potential of +1.25 V *vs*. Ag/AgCl (**Fig. 3C**). The resulting faster sulfur oxidation rate, explains why the optimum-performance period for the BES was the shortest between the thermally activated graphite felt and activated carbon anode treatments (one day in **Table 1**). Adding soluble Fe^2+^ to the thermally activated graphite felt anode with L-cysteine clearly shifted the onset potential to a higher voltage (**Fig. 3C**), reducing the sulfur oxidation rate in our BES, and explaining the longer optimum-performance period (two days in **Table 1**). The no-L-cysteine control (*i*.*e*., no-sulfur control) shifted the onset potential further to a higher voltage, which was anticipated (**Fig. 3C**).

Even though the current densities for the three cyclic voltammograms with activated carbon were approximately four times lower than with thermally activated graphite felt (**Fig. 3C**), we observed the same order of first onset potentials (L-cysteine without soluble Fe^2+^; L-cysteine with soluble Fe^2+^; and without L-cysteine and without soluble Fe^2+^ [control]). We found a mid-point potential of +1.5 V *vs*. Ag/AgCl with the activated carbon with L-cysteine and without soluble Fe^2+^ (**Fig. 1C**). The higher mid-point potential of sulfur oxidation (1.5 *vs*. 1.25 V) and the lower current densities explain why the activated carbon with L-cysteine but without iron showed a longer optimum-performance period for the BES compared to the thermally activated graphite felt (two days *vs*. 1 day in **Table 1**) – the slower sulfur oxidation rate elongated the growth period of methanogens. The addition of soluble Fe^2+^ to the activated carbon with L-cysteine, showed the same result as with thermally activated graphite felt: **(1)** a slower onset potential; and **(2)** a higher mid-point potential (1.6 V *vs*. Ag/AgCl in **Fig. 3C**), explaining the higher optimum-performance period (three days in **Table 1**). Thus, the addition of soluble Fe^2+^, reduced the sulfur oxidation rate.

We suspect that the formation of iron sulfide (FeS) in the electrolyte and/or at the anode slowed sulfur oxidation, allowing the methanogens a longer growth period before the sulfur became limiting. Indeed, the activated carbon-based anodes with soluble Fe^2+^ showed a decreased overall iron concentration throughout time (**Fig. S7**). The initial iron concentration was 0.1 g·L^-1^, continuously decreasing until it stabilized after three days (**Fig. S7**). In addition, we observed the precipitation of FeS, which persisted as a black precipitate. The removal of sulfide through FeS is a desired outcome in water treatment.^34^ Fortunately, it has been shown to benefit *M. thermautotrophicus* ΔH by improving CH_4_ production after adding Fe.^35^ Here, the formation of FeS benefited the microbe by lowering the sulfur oxidation rate. However, FeS is still redox active with sulfur oxidation as an ultimate result, explaining that the maximum optimum-performance period was still only three days (**Table 1**).

### Magnetite, which is an alternative redox mediator to soluble iron, shortened the optimum performance period

The cyclic voltammogram of synthesized magnetite nanoparticles that were immobilized on an Au electrode featured two peaks with a peak splitting of +0.28 V, representing the oxidative peak at -0.12 V *vs*. Ag/AgCl and the reductive peak at -0.41 V *vs*. Ag/AgCl (**Fig. 4A**). The comparison of the standard potential of both magnetite-oxidation reactions with the literature exhibited that the two-electron oxidation of magnetite to maghemite (**Eq. 4**) occurred for our system.^36^ Roberts et al.^37^ had shown that magnetite is prone to develop this irreversible reaction, which would have a negative impact on long-duration BES experiments. Indeed, the cyclic voltammograms of bulk magnetite showed decreasing reaction rates with increasing cycles (**Fig. 4B**), which can be explained by the formation of maghemite (**Eq. 4**).

**Figure 4:**
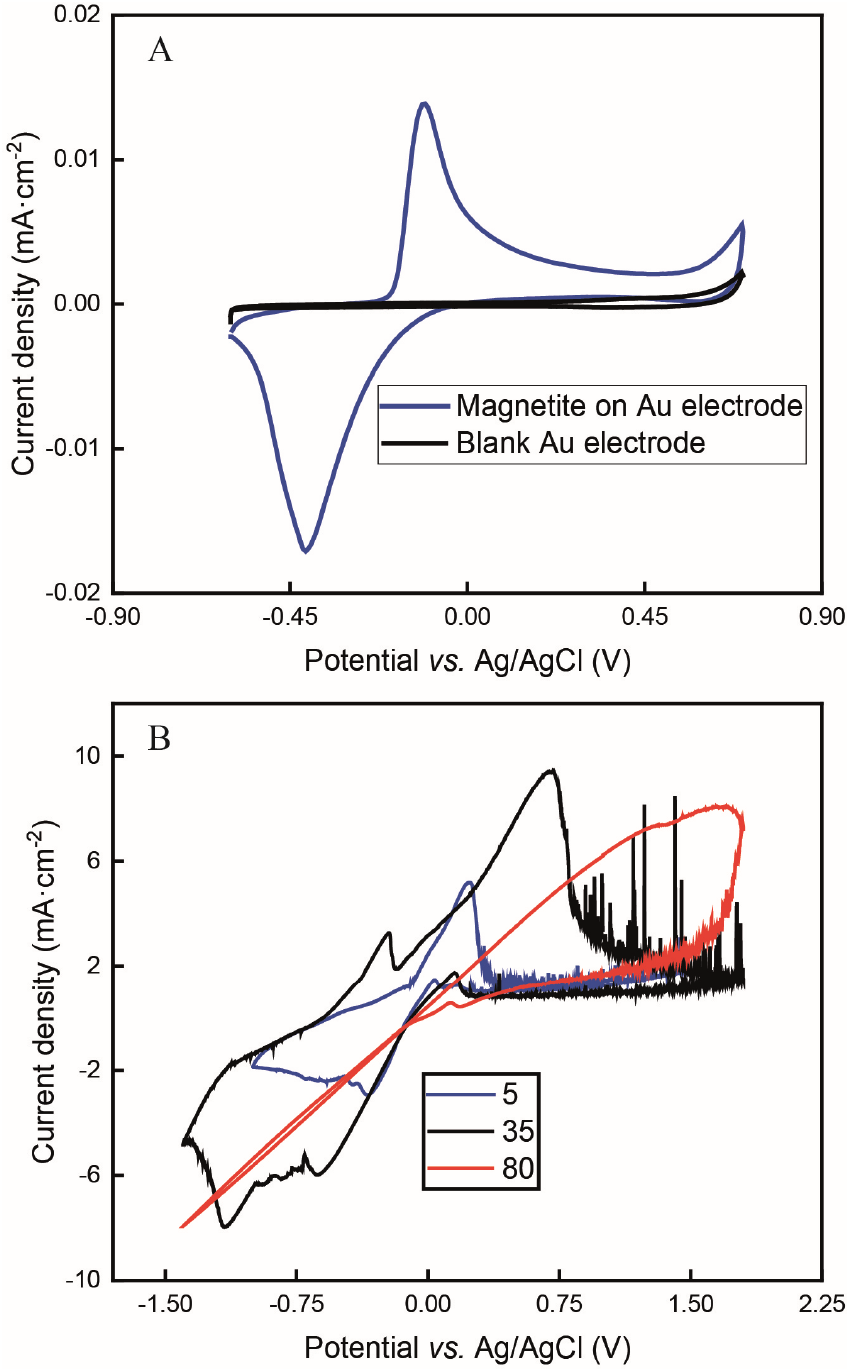
Electrochemical analysis of magnetite: **(A)** cyclic voltammogram of magnetite nanoparticles that were drop-casted on an Au electrode at a scan rate of 20 mVs^-1^ in 66 mM phosphate buffer; and **(B)** cyclic voltammograms of bulk magnetite at cycle 5, 35, and 80 at a scan rate of 20 mVs^-1^ in 50 mM PBS.

To quantify the reaction rates, Tafel slopes were calculated from bulk magnetite during different cycles of the cyclic voltammogram (Supporting Information note 1 and **Fig. 4B**). Bulk magnetite electrodes achieved 187 mV/dec at cycle 5. It transitioned to 617 mV/dec and 1310 mV/dec at cycles 35 and 80, respectively (**Fig. 4B**). The formation of maghemite increased the inner resistance of the anode material to decrease its reaction rate by forming a resistor-type cyclic voltammogram (**Fig. 4B**), which likely would constrain the BES. This is one possible reason why we observed the shorter optimum performance period of two days *vs*. three days for magnetite-coated activated carbon anodes and activated-carbon anodes, respectively. However, more work would be necessary to exclude other possible reasons.

### When sulfur limitations are overcome, carbon at the anode would become limiting next

The oxidation and the removal of the carbon anode itself due to its sacrificing nature, would likely be the next liming factor of our BES due to a lack of accessible carbon to be oxidized. Our proof-of-concept BESs used relatively small geometric anode surfaces of 1.57 cm^2^. For our calculation, we used the ideal reaction of donating four electrons and producing CO_2_ during carbon oxidation (**Eq. 1**). To calculate the radius change (*Δr*) during the two-day optimum performance period of the activated carbon anode (as one example anode), we first calculated the volume (*V*) of carbon that was consumed (**Eqs. 5-8**), which was equivalent to 1.10·10^−2^ cm^3^ (8.95·10^−4^ mol carbon), using the equation of the Coulombic efficiency.^4^ The exposed surface of the activated carbon anode corresponded to a halved cylinder so that the volume after the optimum performance period and its corresponding radius change would be:

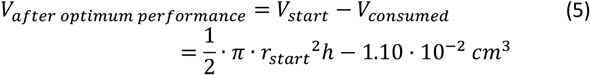

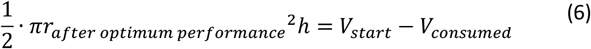

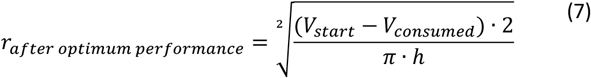

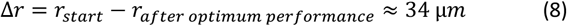

With r_start_ = 0.475 cm and h = 2.175 cm. The relatively small change in the radius of 34 μm shows that the accessible carbon did not limit the system’s performance yet. We could not determine when the carbon would become limiting due to the sulfur limitations. If this occurs, the accessibility of carbon should be increased. This can be achieved by adding more carbon after increasing the ratio of anode surface area to reactor volume or improving the reactivity of the carbon. In addition, anodes could be replaced periodically.

### A final limitation would be the impact of carbon corrosion on the CH_4_ production rates

We observed that the activated carbon-based electrodes started to disintegrate, and small carbon particles were found in suspension that might impact the CH_4_ production rate during long-term operating conditions (considerably longer than three days). We refer to this as carbon corrosion, which could impact the CH_4_ production rate mostly negatively due to: **(1)** the exfoliation of carbon material at the anode, limiting carbon oxidation; and **(2)** the toxicity of a tar-like solution to methanogens. On the other hand, there is a mechanism known from abiotic carbon oxidation systems that could improve CH_4_ production, which we refer to as indirect carbon oxidation by Fe^3+^ (described below). Therefore, we examined carbon corrosion by investigating the surface and the bulk of the anodes with FTIR to track the change of functional groups exhibited at the carbon material. We used the activated-carbon anodes from our abiotic experiments during the measurements of the liquid O_2_ concentration (no soluble iron was added).

Several oxidized functional groups from the initial activated carbon material, such as ketones and phenol groups, were identified by the C=O stretching bond at 1700-1800 cm^-1^ and -OH stretching bonds at 3300-3600 cm^-1^, respectively (**Fig. S8**). The absorbance peak at 2500-3000 cm^-1^ represented the C-H stretching bond of the paraffin oil, which we had added to produce the anodes. The larger absorbance of -OH bonds suggested an alkaline chemical activation of the carbon from the manufacturer. Almost all the -OH stretches disappeared after five days of anode oxidation. The phenol functional group can act as a catalyst for carbon oxidation and facilitate the CO_2_ reaction at the anode. Once the functional group was consumed, the electrode exfoliated in small carbon particles.

While small concentrations of carbon particles in solution might not have a big influence, higher concentrations have been shown to influence methanogenesis. Cavalcante et al.^38^ identified that adding conductive material of carbon particles to a culture of methanogens enhanced methanogenesis. However, inhibition of methanogenesis occurred even at lower carbon material concentrations when the inoculation was performed at a relatively low cell density. In addition, Umetsu et al.^39^ showed that *M. thermautotrophicus* ΔH adsorbed on -OH and -COOH functional groups and prevented hydrogen or mineral uptake, inhibiting methanogenesis.

We also found that carbon corrosion and the reaction of the released carbon with Fe^3+^ (**Eq. 9**) could explain another stimulation of the CH_4_ production rate besides: **(1)** reducing the sulfur oxidation rate due to FeS formation; and **(2)** the presence of conductive particles. Iron would need to be present in our membrane-less BES for this other stimulation, and we refer to this mechanism as indirect carbon oxidation by Fe^3+^. The continuous addition of soluble Fe^2+^ stimulated abiotic carbon oxidation systems because Fe^2+^ is directly reduced at the carbon-based anode to Fe^3+^ (**Eq. 3**). However, even without continuous soluble Fe^2+^ addition, a current increase can be anticipated due to the circular nature of the electron-mediating process, with the ultimate electron donor coming from the carbon-based anode material than the continuously added Fe^2+^. This is due to indirect carbon oxidation by Fe^3+^ using carbon in the solution that originated from carbon corrosion at the anode (**Fig. 5**). Indirect carbon oxidation reduces Fe^3+^ back to Fe^2+^ in the solution, while carbon is oxidized to CO_2_ (**Eq. 9** and **Fig. 5**). The reactions of the electron-mediating process can be described as follows:

**Figure 5:**
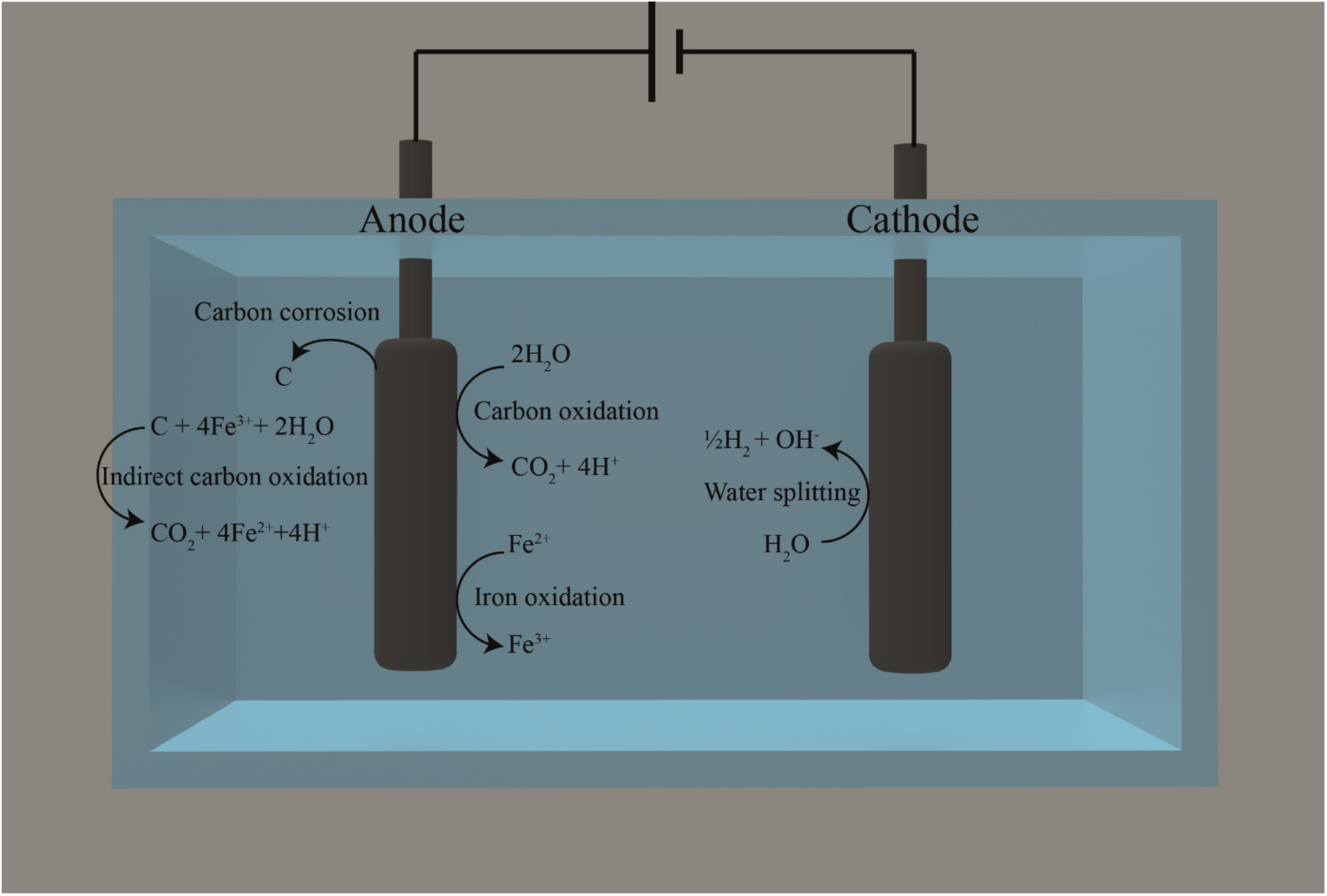
Overview of the reactions that occur at the anode and cathode during the carbon oxidation using thermally activated graphite felt and activated carbon anodes with 0.1 g·L^-1^ of Fe^2+^. Water splitting takes place at the cathode to produce H_2_. The main reaction at the anode is carbon oxidation, followed by the oxidation of Fe^2+^ and indirect carbon oxidation by the formed Fe^3+^. For more information, see section on the impact of carbon corrosion on the CH_4_ production rates.

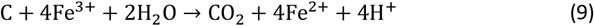

Because the reduction and oxidation of soluble iron species occur at the anode and with carbon in solution in the electrolyte, respectively, there is a continuous reformation of Fe^2+^ to maintain a higher current density for the electrochemical cell. An advantage of the circular nature of this electron-mediating process is that Fe^3+^ is oxidized in the electrolyte with carbon and not at the cathode, which, as a competing process, would have reduced the Coulombic efficiency for methane production.

We do not know whether carbon corrosion would have positive or negative effects on the CH_4_ production rate and the duration of the optimum-performance period for our BES. We would first need to solve the sulfur oxidation problem. Likely, there may be an initial positive effect when carbon corrosion is only starting to occur, but when more and more carbon particles accumulate, we may observe a negative effect of carbon corrosion. Therefore, measures should be taken to reduce the negative effects of carbon corrosion on the anode activity and microbial activity to lengthen the optimum performance period of anode oxidation with sacrificial anodes.

### Carbon oxidation with sacrificial anodes for single-chamber BES is ready for research purposes within the field of microbial electrosynthesis

We showed that implementing carbon oxidation with sacrificial anodes in a single-chamber BES can perform microbial electrosynthesis. We cultivated *M. thermautotrophicus* ΔH with *in-situ* produced H_2_ at the cathode under constant pH and temperature conditions to produce CH_4_ for an optimum performance period of three days. The growth behavior followed a similar trend for all BESs until the amount of sulfur limited the CH_4_ production rates. Our maximum cathode-based geometric CH_4_ production rate was already compatible with other studies due to similarly set current densities.^4^ Further optimizations of our system should not only increase the geometric CH_4_ production rate but also prolong the production days to several weeks. With the relatively low current densities of microbial electrosynthesis, our system is already useful for research purposes because the sulfur will not become limiting that fast with lower current densities. Our work can also inform the earlier studies with a single-chamber BES in the field of perchlorate removal.^21^ We advise the authors to replace the graphite anode materials with activated-carbon anode materials and to add soluble iron to increase the performance.

### Before the industrial application of carbon oxidation with sacrificial anodes in BES for microbial electrosynthesis can commence, several breakthroughs are required

Our system would have a considerably lower theoretical power consumption than membrane-based two-compartment cells due to the lower potential to operate the BES (**Fig. S4** and **S9**). After optimizing the single-chamber BES and making it continuous, a direct comparison between the production rates and power consumption rates should be performed with a conventional membrane-based cell for microbial electrosynthesis. Another advantage of the single-chamber cell compared to a membrane-based cell would be the lower capital costs due to the absence of ion-exchange membranes, albeit a full techno-economic analysis would be necessary to be conclusive. These advantages make our system a desirable opportunity for scale-up. However, several problems must be overcome.

We must prolong the optimum performance period to several months rather than a few days before scale-up can commence. We observed the limiting factor of sulfur availability in our membrane-less BES, which had a promising current density. Utilizing a microbe that is not reliant on a redox-active sulfur compound as either a sulfur source for growth or to provide reducing power would be one route to overcome the utilization of a single-chamber BES. If this problem can be overcome, we predict two additional limitations for our membrane-less BES: **(1)** the availability of accessible carbon; and **(2)** carbon corrosion. Because carbon gets consumed over time, innovative BESs must be designed to refill the consumed carbon at the anode.

Undoubtedly, carbon corrosion would inhibit the growth of microbial cells during operating periods of several weeks, depending on the current densities. In addition, carbon corrosion would introduce a higher content of -OH, -C=O, and -COOH functional groups, increasing the charge transfer resistance of the carbon material and decreasing the electrochemical performance.^40^ Consequently, the carbon anode decreases its conductivity and current density, while deteriorating. Ideally, carbon corrosion is avoided by efficiently transforming carbon into CO_2_ or CO. Studies to mitigate carbon corrosion currently tend to use either expensive catalysts to protect the carbon material or structurally stronger carbon, such as graphene or carbon nanotubes.^41^ In our case, excessive carbon corrosion has to be avoided, and a continuous system rather than our batch system could remove the by-products. Furthermore, mesoporous carbon would enlarge the surface area and the porosity, allowing more reaction sites for carbon oxidation rather than carbon corrosion.

## CONCLUSION

We have proven the concept that microbial electrosynthesis is possible in a BES without membranes by utilizing carbon oxidation with sacrificial anodes. Our BES produced CH_4_ with an oxygen-sensitive thermophilic methanogen at a cathode-based geometric current density of 1 mA cm^-2^ for 3 days after which sulfur became limiting in the growth medium due to sulfur oxidation at the anode. The system is ready for microbial electrosynthesis experiments with relatively low current densities to perform basic research. However, for industrial applications, several breakthroughs are necessary to overcome limitations at the anode, such as sulfur oxidation, refilling of carbon, and protecting against carbon corrosion.

## EXPERIMENTAL SECTION

### Chemicals, strain, and medium

All chemicals were used as received unless otherwise noted in the text. These chemicals were of technical or analytical grade and were purchased from Sigma-Aldrich (St. Louis, USA), Alfa Aeser (Haverhill, USA), Merck (Darmstadt, Germany), Thermo-Fischer (Waltham, USA), VWR (Radnor, USA), and Carl Roth (Karlsruhe, Germany). The strain *M. thermautotrophicus Δ*H (DSM 1053) was obtained from the DSMZ (Braunschweig, Germany). A pre-culture was grown at 60°C in 100-mL serum bottles with 20 mL of modified anaerobic MS-Medium^31^ containing (per liter): 6.77 g NaHCO_3_; 0.17 g K_2_HPO_4_; 0.235 g (NH_4_)_2_SO_4_; 0.09 g MgSO_4_; 0.06 g CaCl_2_ × 2 H_2_O; 0.23 g KH_2_PO_4_; 1 mL trace elements (10x); 1 mL of 4 M H_2_SO_4_; 0.5 g L-cysteine hydrochloride. The trace-element solution was a 10x stock solution containing (per 100 mL): 0.2 g nitrilotriacetic acid; 3 g MgSO_4_ x 7 H_2_O; 0.5 g MnSO_4_; 0.3 g FeSO_4_ x 7 H_2_O; 0.11 g CoSO_4_ x 7 H_2_O; 0.068 g CaCO_3_; 0.18 g ZnSO_4_; 0.01 g CuSO_4_ x 5 H_2_O; 0.018 g KAl(SO_4_)_2_ x 12 H_2_O; 0.01 g H_3_BO_3_; 0.01 g Na_2_MoO_4_ x 2 H_2_O; 0.48 g (NH_4_)_2_Ni(SO_4_)_2_ x 6 H_2_O; 0.01 g Na_2_WO_4_ x 2 H_2_O; 0.01 g Na_2_SeO_4_. The trace-element solution was adjusted to a pH value of 1 with H_2_SO_4_. The MS-Medium was buffered only in the bioelectrochemical system by adding 2.5 M 3-(*N*-morpholino)propanesulfonic acid (MOPS) at pH = 7.2 for a final media concentration of 25 mM. 25mM Ti-NTA was prepared according to Moench et al.^42^. All solutions were sterilized either through a sterile filter or autoclaved and made anaerobic by sparging with N_2_ or N_2_/CO_2_.

### Preparation of carbon-based electrodes

The activated carbon served not only as active material but also as a conductive and surface enlarging material and was used as purchased (∼100 mesh particle size (∼150 μm), DARCO^®^, Cabot Norit Americas Inc, Marshall, USA). The activated carbon and paraffin oil (Sigma-Aldrich) was homogeneously mixed in a mortar at 70%:30% to form a paste ^43^. The resulting paste was pelleted into a self-made electrode holder and cured in an 80°C vacuum oven for 4 h.

Activated-carbon-associated magnetite was synthesized through a co-precipitation method by mixing activated carbon with 2 M FeCl_3_ and 1 M FeCl_2_.^44^ Then, 20 % NH_4_OH was slowly dropped into the solution until pH 9. The solution was stirred for 1 h to complete the synthesis of magnetite. The precipitate was centrifuged, cleaned with deionized water until pH 7, and dried in a vacuum oven at 80°C for 4 h. Different amounts of activated carbon to magnetite were used during the synthesis ranging from 0% to 60% of activated carbon. In this study, activated carbon that was co-precipitated with magnetite is called magnetite-coated activated carbon (**Supporting Information**). The 60% activated carbon and 40 % maghemite mixture was synthesized by adding 60 % activated carbon and 40 % magnetite mixture into an aerated oven at 80°C for 24 h.

### Measurement of dissolved O_2_

The O_2_ measurements were carried out in a self-designed five-neck round flask. To measure the dissolved O_2_, an O_2_-sensitive foil (0-100% O_2_, PreSens Precision sensing, Regensburg, Germany) was glued at the mid-height of the flask. The electrochemical cell was operated as a two-electrode setup with a 1x1-cm carbon cloth electrode as the cathode. A platinum wire, a graphite rod, a thermally activated graphite felt, a carbon cloth, activated carbon, and activated carbon that was coated with magnetite were used as anodes. Thermally activated graphite felt and activated carbon anodes were used with addition of soluble iron. The measurements were carried out in 0.066 M phosphate buffer at room temperature and 60°C. The phosphate solution was buffered at 7.2 to mimic the pH of the MS medium used in the bioelectrochemical system. Before each experiment, the cells were purged with N_2_ gas for 1 h to secure an O_2_-free solution.

### Sample analyses

The bioelectrochemical systems were sampled once per day. The gas compositions of the bioelectrochemical systems were sampled with a 500-μL gastight syringe (Hamilton, Reno, USA) and analyzed using a gas chromatograph (SRI GC, Torrance, USA). The gas chromatograph used a Haysep D column (length 3m, outer diameter 1/8”, SRI GC) and a Molsieve 13X column (length 3m, outer diameter 1/8”, SRI GC). It was equipped with a thermal coupled detector and a flame ionization detector to measure N_2_, O_2_, H_2,_ and CO_2_, and CH_4_, respectively. N_2_ and H_2_ were used as carrier gas. Coulombic efficiencies for CH_4_ were calculated according to Kracke et al.^4^ Cell growth was measured by cell counting with a light microscope (X41, Olympus, Shinjuku, Japan). The anodes were collected after the experiment and stored anoxically at -20°C for further analyses. The outer layer and the bulk of magnetite particles were analyzed through a sequential extraction. First, the outer layer of magnetite particles was dissolved by acidifying with 1 M HCl for 1 h. It was followed by a complete dissolution of the magnetite particle by acidifying with 6 M HCl for 24 h. The iron content from the liquid samples and the oxidation state of iron in the magnetite electrodes were further analyzed *via* the Ferrozine assay.^44, 45^ Sulfide concentrations were determined photometrically based on a modified version of Cline and Gilboa-Garber ^46, 47^. Sulfide samples were stabilized with Zn(ac)_2_ and stored at -20°C. The diamine reagent (N,N-Dimethyl-p-phenylenediamine sulfate salt and FeCl_3_) was mixed with the stabilized sample at a 1:1 ratio and kept in the dark for 30 min for the reagents to react. The samples were measured at 565 nm. The ORP was measured with an ORP electrode (Hanna Instruments, Vöhringen, Germany) connected to an ORP controller (Alpha pH 800, Eutech Instruments, Singapore).

### Electrochemical system

All electrochemical measurements (cyclic voltammetry, linear sweep voltammetry, and chronopotentiometry) were conducted in a five-neck round flask with a working volume of 50 mL. The electrochemical cells were operated as a two-electrode or a three-electrode setup and controlled *via* a potentiostat (VMP3, Bio-Logic, Grenoble, France). An Ag/AgCl (3M KCl, 0.210 V *vs*. SHE) in-house-made reference electrode was used for the three-electrode setup. All measurements with a reference electrode were ohmic-drop corrected and were determined by potentiostatic electrochemical-impedance spectroscopy. Ohmic-drop-corrected potentials were calculated following the method of Sun et al.^48^ The reaction kinetics of carbon and magnetite were calculated with the Butler-Volmer equation, using linear sweep voltammetry and cyclic voltammetry measurements (Supporting Information). The 1x1-cm carbon cloth electrode (PANEX 30-PW06, Zoltek, St. Louis, USA), thermally activated graphite felt (Sigracell, SGL carbon, Bonn, Germany) and the platinum wire (Bioanalytical Systems, Inc., West Lafayette, USA) were used as cathodes, and the carbon-based electrode, graphite rod (Mersen, Suhl, Germany), carbon cloth (Zoltek), thermally activated graphite felt (SGL carbon), the carbon paste electrode holder (Bio-Logic), and the gold electrode (Bio-Logic) were used as anodes depending on their intended application, which is described in the results part. A titanium wire (Sigma Aldrich, St. Louis, USA) was threaded through the carbon cloth and thermally activated graphite felt, and glued with carbon conductive cement (Thermo-Fischer, Waltham, USA) for better conductivity. The temperature was either room temperature or controlled by a water bath at 60°C (ICC basic IB R RO 15 eco, IKA, Staufen im Breisgau, Germany). The supporting electrolyte for the abiotic experiments was phosphate buffer (0.066 M) or phosphate buffered saline (0.05 M), with the pH buffered at 7.2. Before each abiotic electrochemical experiment, a high-purity N_2_ gas was introduced to remove O_2_. The bioelectrochemical systems were either autoclaved or chemically sterilized before the beginning of each experiment. The reactors were filled with 50-mL MS-Medium, were kept anaerobic by sparging with 80:20 (*v/v*) N_2_/CO_2_ through a sterile filter, and were continuously stirred with a magnetic stir bar. The reactors were operated in batch mode and were fed for 5 min with pure CO_2_ as the sole carbon source *prior* to the experiment and every day for 1 min after liquid sampling to exchange the headspace. After feeding, an overpressure of 80 mbar of CO_2_ was kept in the headspace. The electrochemical cells were submerged in a water bath to prevent heat loss for optimal growth of *M. thermautotrophicus* ΔH at 60°C.

### Immobilization of magnetite

Magnetite was immobilized by dispersing magnetite nanoparticles onto an Au electrode (Biologic) to test its electrochemical behavior. The protocol is an adjusted version from Sun et al.^48^ Briefly, ten milligrams of the active powder and 40 μL of Nafion were mixed with 960 μL 70% ethanol and sonicated for 15 min. Then, 10 μL were drop casted onto the surface of the Au electrode and dried at room temperature. To avoid iron from oxidizing, the preparations for the drop casted magnetite were performed inside a glovebox (mBraun, Garching, Germany).

### SEM, EDX, X-ray diffraction, and FTIR

A 50-μL aliquot of 0.1% poly-L-lysine solution was placed onto a glass slide and was dried for 1 h in a 60°C incubator. The samples were placed onto a poly-L-lysine coated glass slide that was attached to aluminum stubs, using carbon adhesive tabs. The sample was coated with a 10-nm deposition of platinum, using a BAL-TEC™ SCD 005 sputter coater (Leica biosystems, Nussloch, Germany). The purpose of the coating was to reduce any charging effects during scanning electron microscope (SEM) analysis. All images were taken in secondary-electron (SE) mode using a Zeiss Crossbeam 550L focused ion beam (FIB) SEM (Zeiss, Wetzlar, Germany), which was operated with an acceleration voltage of 2 kV. The SEM was equipped with an Oxford Instrument energy dispersive spectrometry detector (Ultim Max, Oxford Instrument, Abingdon, United Kingdom). Elemental maps of iron and carbon were obtained along with corresponding spot analysis.

X-ray diffraction (XRD) was carried out on dry material using Bruker’s D8 Discover GADDS XRD2 micro-diffractometer that was equipped with a Co-anode (Cu Kα radiation, λ = 0.154 nm) at parameters of 30 kV/30mA (Billerica, USA). The total measurement time was 240 s at two detector positions, which were 15° and 40°. Fourier transform infrared (FTIR) spectroscopy was performed on a Vertex 80v equipped with an RT-DLaTGS detector (Bruker, Billerica, USA). 0.5 mg of sample was mixed homogenously with 250 mg KBr (FTIR grade, Carl Roth, Karlsruhe, Germany) and compressed at 10T to form a pellet. Samples were scanned 32 times in the range of 400-4500 cm_-1_.

## Author Contributions

L.T.A., T.S., and R.B.-G. initiated the work. N.R., T.S., and L.T.A. designed the experiments. N.R. performed the laboratory experiments and analyzed the data. L.T.A. and T.S. supervised the project. N.R. and L.T.A. wrote the manuscript. All edited the paper and approved the final version.

## Acknowledgements

This work was supported by the Deutsche Forschungsgemeinschaft (DFG, German Research Foundation, EBiotech, SPP2240; L.T.A.) – project number 445506379, the Alexander von Humboldt Foundation in the framework of the Alexander von Humboldt Professorship (L.T.A.), and The Novo Nordisk Foundation CO_2_ Research Center with grant number NNF21SA0072700 (L.T.A.). L.T.A and A.K. are also grateful to funding from the Deutsche Forschungsgemeinschaft under Germany’
ss Excellence Strategy – EXC 2124 – 390838134. R.B-G. was supported by an FPI grant (BES-2015-074229) from the Spanish Ministry of Economy and Competitiveness (MINECO) within the research project CTQ2014-53718R. The authors gratefully acknowledge the Tübingen Structural Microscopy Core Facility (Funded by the Federal Ministry of Education and Research (BMBF) and the Baden-Württemberg Ministry of Science as part of the Excellence Strategy of the German Federal and State Governments) for their support and assistance in this work. The authors also thank the German Research Foundation DFG (INST 37/1027-1 FUGG) for the financial support provided for the acquisition of the cryogenic focused ion beam scanning electron microscope. We thank Prof. Dr. Uwe Schröder (University of Greifswald) for helpful and insightful discussions.

