## Supplementary material for "Carbon oxidation with sacrificial anodes to inhibit O_2_ evolution in membrane-less bioelectrochemical systems for microbial electrosynthesis": SI

**Supplementary note 1:**

**Determination of the exchange current density and Tafel slope**

For the determination of the exchange current density for the carbon oxidation, OER, and HER, the Butler-Volmer equation (**Eq. S1**) was used and transformed to determine $j_0$ from its interception at the y-axis.[1] The Butler-Volmer equation (**Eq. S1**) is described as followed:

$$j = j_0 \left\{ exp\left[\frac{\alpha_a zF}{RT}\eta\right] - exp\left[\frac{\alpha_c zF}{RT}\eta\right]\right\} \tag{S1}$$

With j the current density, $j_0$ the exchange current density, $\alpha_a$ and $\alpha_c$ the anodic and cathodic charge transfer coefficient, respectively, z the number of electrons involved in the reaction, F the Faraday constant, R the universal gas constant, T the temperature, and η the activation overpotential.

When η>>0, the **Eq. S1** simplifies to **Eq. S2** :

$$\eta = \frac{2.3\ RT}{\alpha F} log j_0 - \frac{2.3\ RT}{\alpha F} log j \tag{S2}$$

**Eq. S2** can be transformed into the Tafel equation to calculate the Tafel slope, which was used to calculate the Tafel slopes of magnetite:

$$\eta = \pm \frac{2.3\ RT}{\alpha F} log \frac{j}{j_0} + A \tag{S3}$$

$$\eta = B log \frac{j}{j_0} + A \tag{S4}$$

With B the Tafel slope, and A the symmetry factor. The exchange current density $j_0$ and the Tafel slope are calculated from log|j| *vs.* η by the interception of the y-axis and the slope, respectively.

**Supplementary note 2:**

**Magnetite analyses**

We investigated magnetite as a possible redox mediator. We fabricated magnetite anodes with activated carbon, which is our carbon source for carbon oxidation, to increase the conductivity of magnetite, which is a semi-conductor.[2] Since the conductivity of the electrode can vary depending on the amount of activated carbon used, we investigated which ratio of magnetite to activated carbon is suitable for higher current generation, longevity, and inhibition of $O_2$ evolution. Therefore, we performed abiotic tests with different ratios of magnetite and activated carbon ranging from 20% to 100% of the total activated carbon. We used the chronoamperometric method to determine the electronic conductivity of our mixtures and set the WE at +0.6 V *vs*. Ag/AgCl. The current of a chronoamperometric graph decreased exponentially to reach an asymptotic value, which was used to calculate the conductivity (**Fig. S10**). The specific electronic conductivity $\sigma$ was calculated (**Eq. S5**).

$$\sigma = \frac{I \cdot D}{E \cdot A} \tag{S5}$$

Where $I$ is the asymptotic current, $D$ is the thickness of the electrode ($D$ = 0.4 cm), $E$ is the applied potential, and $A$ is the surface area of the electrode ($A$ = 0.28 cm$^2$). The calculated specific electronic conductivities of 40% magnetite and 60% activated carbon showed the highest electronic conductivity with $3.89 \times 10^{-4}$ S·cm$^{-1}$ and was further analyzed for its ability to inhibit $O_2$ (**Table S2**).

A critical aspect of sufficient electrical conductivity is a homogenous distribution of magnetite on activated carbon particles. Ideally, only a thin layer of magnetite is adsorbed on the surface of the activated carbon. To analyze the distribution of magnetite on activated carbon, we characterized synthesized magnetite on activated carbon with X-ray diffraction (XRD). We compared synthesized magnetite with the following materials: **(1)** pure magnetite; **(2)** pure activated carbon; and **(3)** commercial magnetite (**Fig. S11**). The XRD patterns of magnetite, magnetite on activated carbon, and commercial magnetite show overlapping diffraction peaks for the whole spectrum, confirming that our synthesis contains magnetite. Further comparison of the XRD peaks shows a good agreement with standard patterns, indicating high crystalline quality. However, because maghemite's XRD pattern is virtually indistinguishable from the XRD pattern of magnetite, we analyzed the Fe(II)/Fe(III) ratio with the ferrozine assay. The ratio of Fe(II)/Fe(III) found in our samples corresponded to 0.40 ± 0.1, which is 20% lower than the ideal ratio of 0.5 for magnetite. This means that 20% of our material was oxidized, likely to maghemite. The XRD pattern of the activated carbon can be attributed to amorphous carbon that is also observed in magnetite coated activated carbon.

Scanning electron micrograph of magnetite on activated carbon demonstrated a rough and porous particle (**Fig. S12A**). Furthermore, close-up views of the particle showed that magnetite was synthesized as round-shaped nanoparticles (**Fig. S12B,C**). However, it was not possible to identify whether magnetite was homogeneously coated on activated carbon. To underline the homogenous distribution of magnetite, we performed EDX to detect the iron on the surface. By detecting the Fe with Fe Kα energy-dispersive X-ray spectroscopy (EDX) elemental mapping of magnetite-coated activated carbon, we detected the distribution of Fe. The overlay of a magnetite-coated activated carbon particle SEM micrograph with the elemental mapping of Fe confirmed that Fe was very well dispersed on top of the activated carbon particle (**Fig. S13** and **S14**).

**The stability of magnetite anodes is primarily determined by the crystal structure and purity of the magnetite mineral**

We calculated the conductivity of different magnetite to activated carbon ratios and found that the ratio of 40%:60% magnetite to activated carbon obtained higher conductivities for our electrodes (**Eq. 5** and **Fig. S10**). To understand how this ratio functions in stability and longevity, we prepared triplicates of 40%:60% magnetite to activated carbon electrodes as our counter electrodes and poised the carbon cloth WE at -0.8 V *vs.* Ag/AgCl. After 7 days, the liquid/electrode interface developed an orange precipitate of Fe(III) at each electrode. After the experiment, we analyzed each Fe(II)/Fe(III) electrode ratio with a sequential extraction. We compared them with the starting material (**Fig. S15**) and analyzed the abundance of Fe(II) and Fe(III) for each section (electrode liquid interface, middle section, upper section). Here, the starting material was partially oxidized with a stoichiometric ratio of around 0.44. After 7 days of oxidation, the outer layer of the magnetite particle was nearly completed oxidized, while the stoichiometric ratio of the bulk was similar to the initial stoichiometric ratio of magnetite. In conclusion, activated carbon coated magnetite showed nearly complete oxidation at the outer layer that might prevent a stable and durable anode. Refer to the main text for further informations on the durability of magnetite anodes under section “Magnetite, which is an alternative redox mediator to soluble iron, shortened the optimum performance period”

**Table S1│ Comparison of abiotic and biotic current densities with their respective coulombic efficiencies and references**

| Product | Current density [mA $cm^{-2}$] | Coulombic Efficiency [%] |
|---|---|---|
| | **Abiotic** | |
| $H_2$ | Up to 2000 | 99 [3] |
| Ethanol | 250 | 46 [4] |
| $CH_4$ | 200 | 70 [5] |
| | **Biotic** | |
| Acetate | 10 | 100 [6] |
| $CH_4$ | 1 | 99 [7] |

**Table S2│The electronic conductivity of different activated carbon to magnetite and hematite percentages calculated by chronoamperometry (CA) method.**

| Activated carbon to magnetite | Electronic conductivity [S/cm] |
|---|---|
| 100% activated carbon | $5.08 \times 10^{-8}$ |
| 60% activated carbon, 40% magnetite | $3.89 \times 10^{-4}$ |
| 40% activated carbon, 60% magnetite | $2.91 \times 10^{-4}$ |
| 20% activated carbon, 80% magnetite | $1.91 \times 10^{-4}$ |
| 60% activated carbon, 40% maghemite | $5.31 \times 10^{-6}$ |

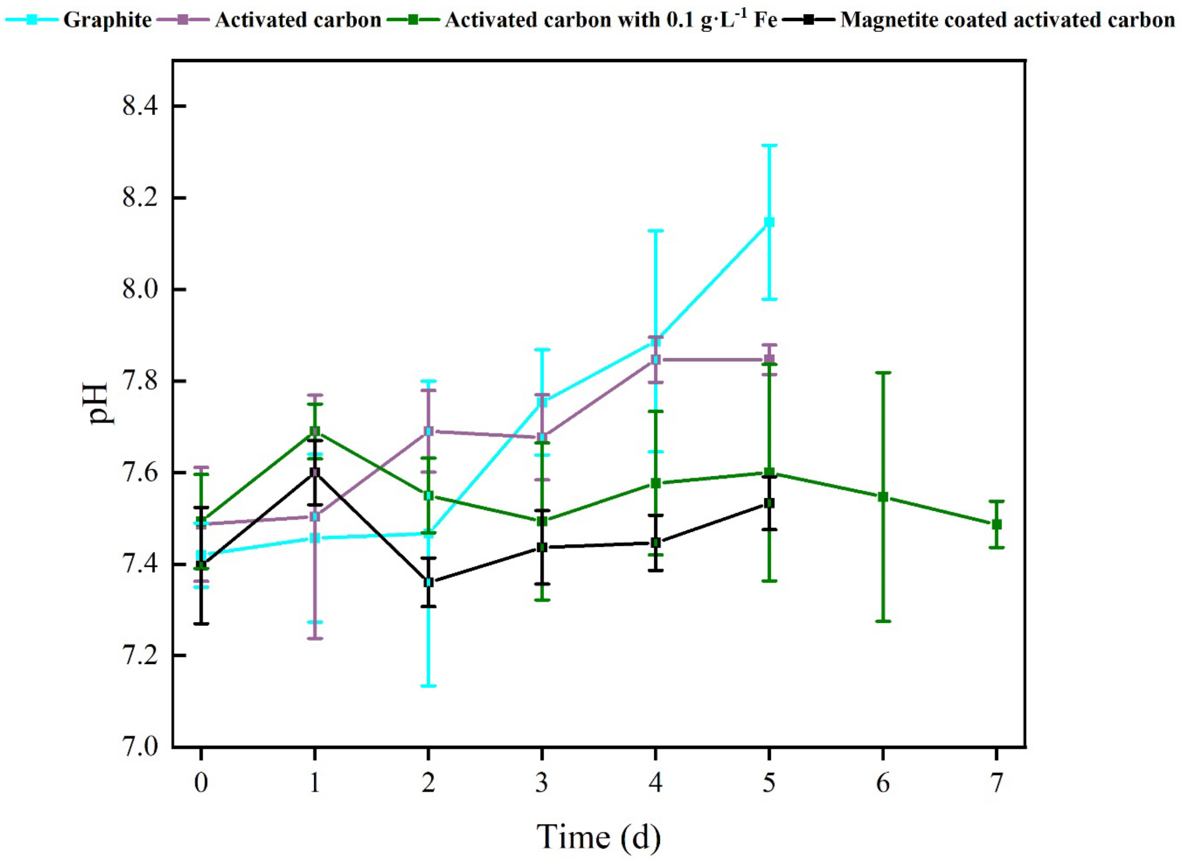


**Figure S1:** Measured pH during some of the BES experiments using graphite, activated carbon with and without added iron, and magnetite-coated activated-carbon anodes.

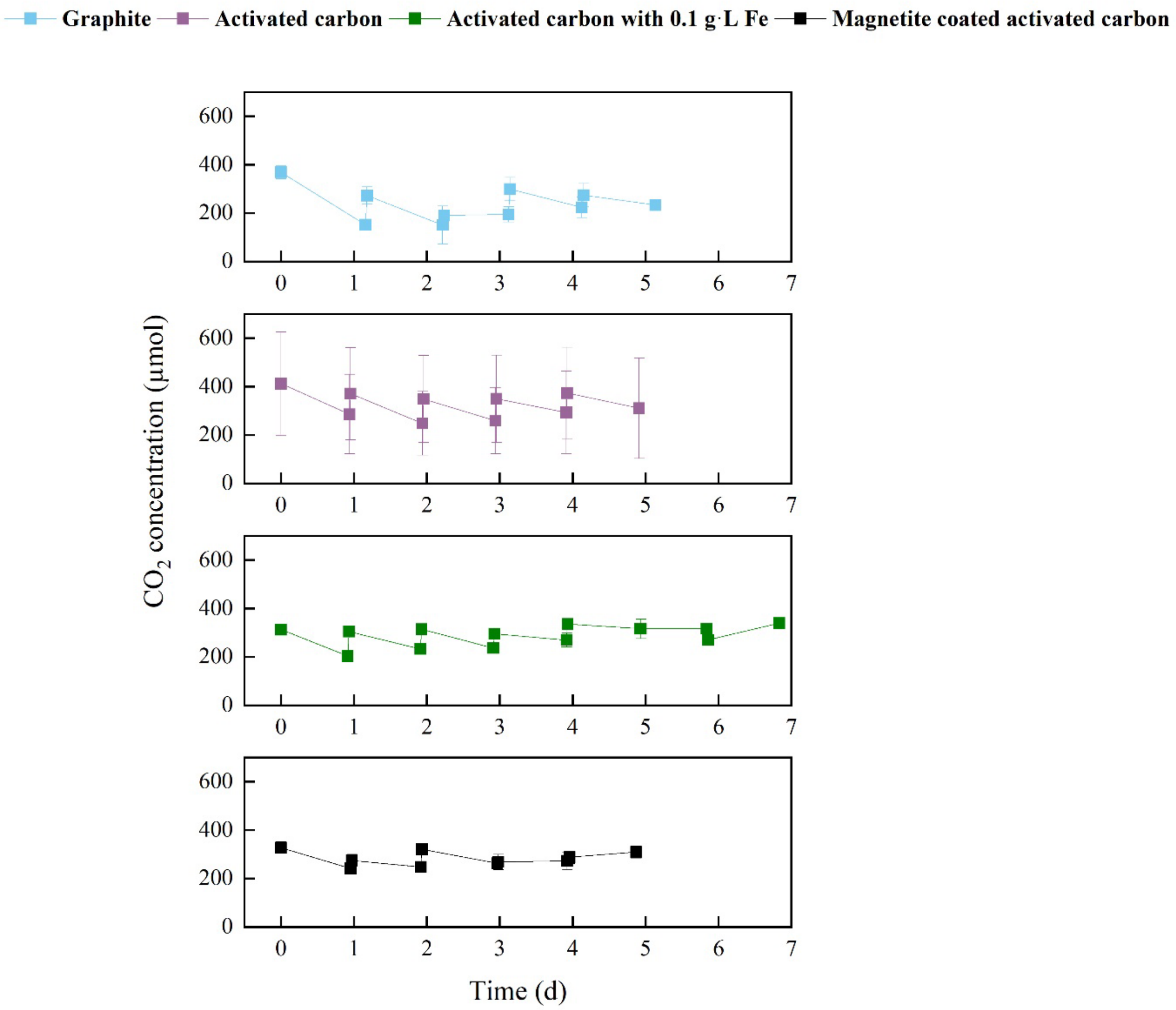


**Figure S2:** $CO_2$ concentration during batch experiments throughout time with the thermophilic archaeon *M. thermautotrophicus* ΔH. Graphite-rod and activated-carbon anodes were included with three treatments for activated carbon, including iron additions (soluble iron and magnetite). The error bars show the standard deviation for the average data from triplicate systems.

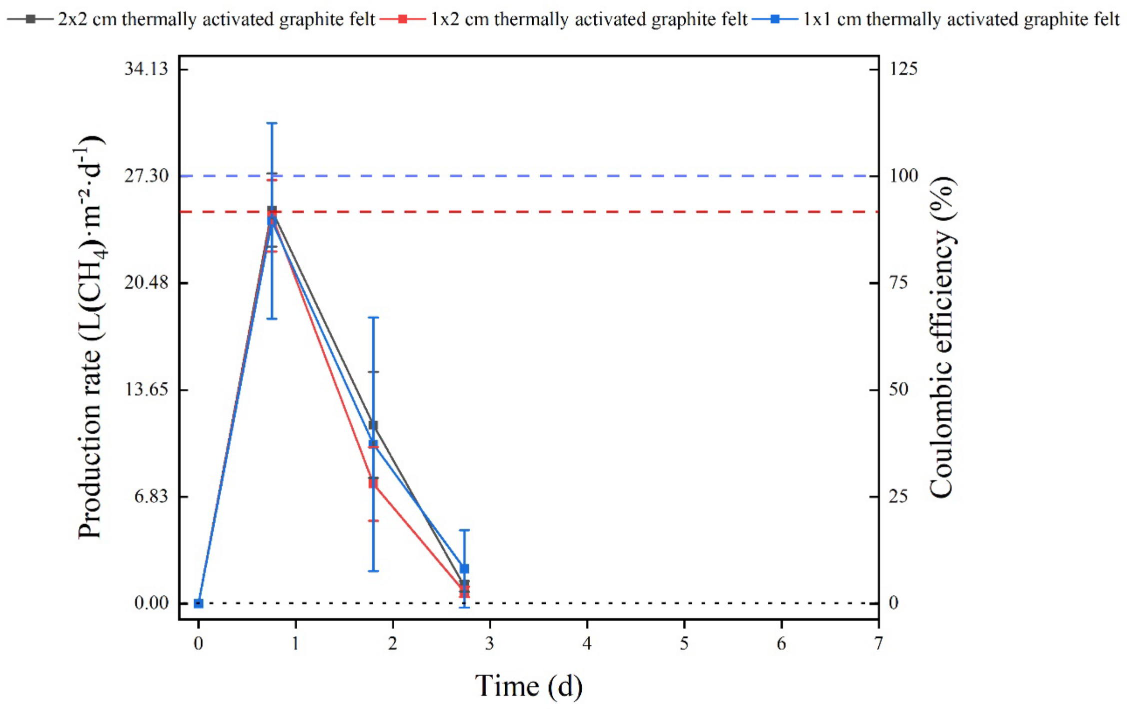


**Figure S3:** $CH_4$ production rates corrected to the cathode surface area and Coulombic efficiencies (at I = 1 $A \cdot cm^{-2}$) during batch experiments in our BES throughout time with L-cysteine and with the thermophilic archaeon *M. thermautotrophicus* ΔH. Thermally activated graphite felt anodes were included with three different conditions (1x1 cm , 1x2 cm graphite felt, and 2x2 cm). The error bars show the standard deviation for the average data from triplicate systems. Carbon cloth was used as the cathode: **(**Y-axis 1) cathode-based geometric $CH_4$ production rates; and **(**Y-axis 2**)** Coulombic efficiencies.

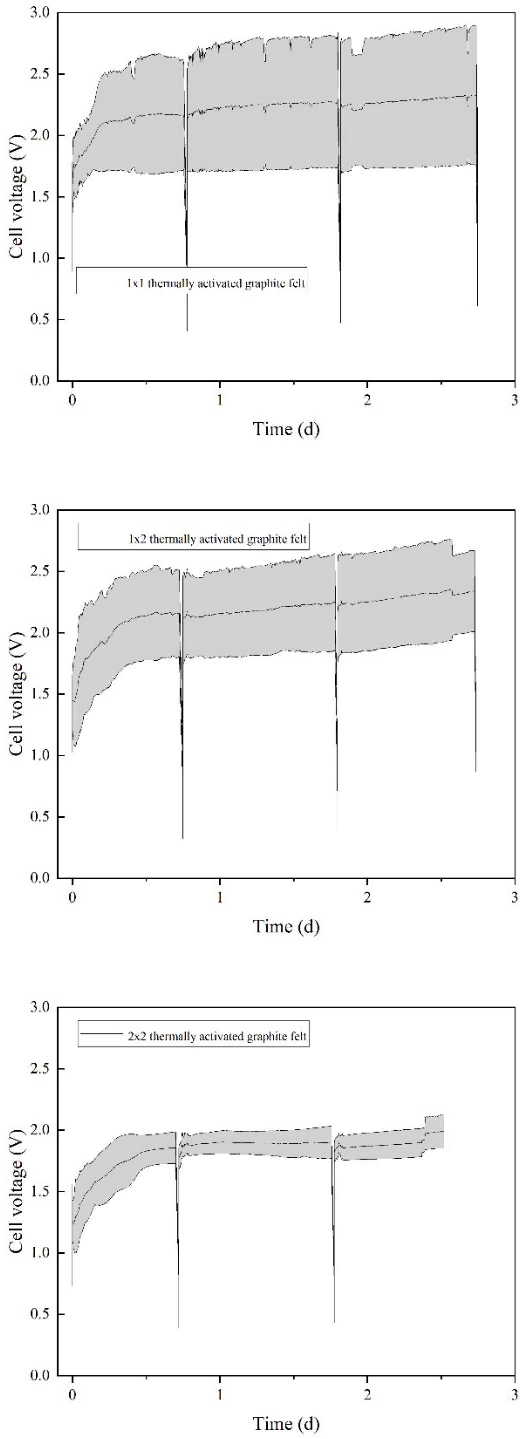


**Figure S4:** Cell voltages of the BES experiment performed in triplicates using three different sizes (1x1, 1x2, and 2x2 cm) of the thermally activated graphite felt as anode.

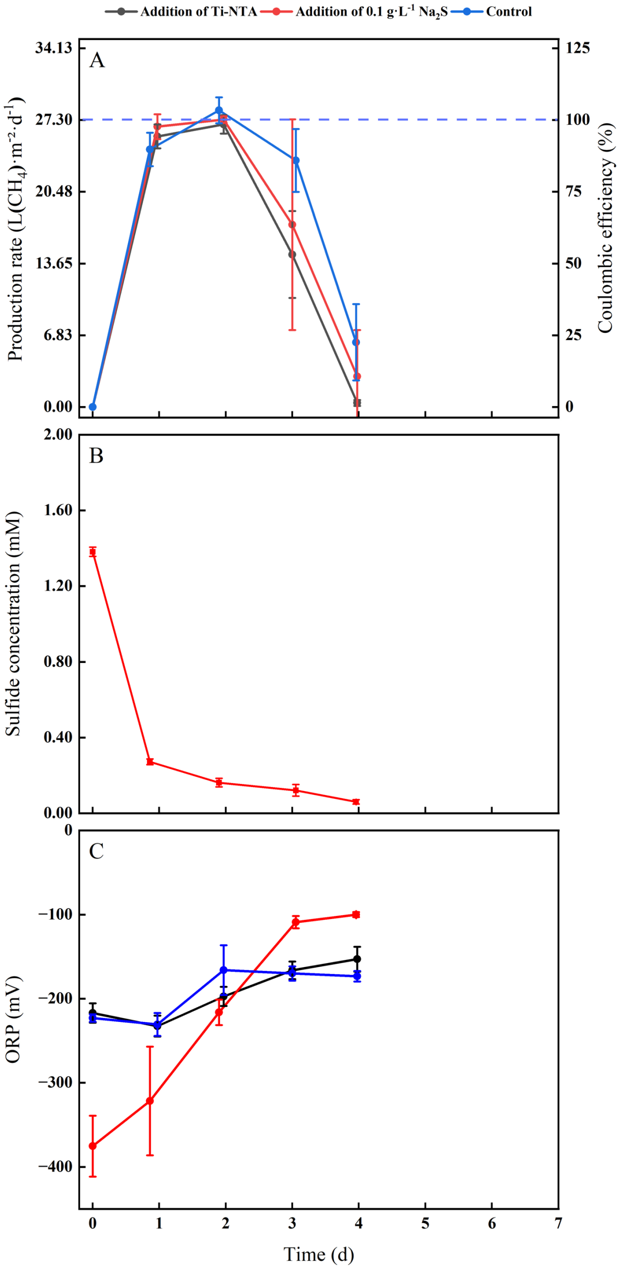


**Figure S5:** BES experiments with different reducing agents: (**A**) $CH_4$ production rates corrected to the cathode surface area and Coulombic efficiencies (at I = 1 A·cm$^{-2}$) during batch experiments throughout time with the thermophilic archaeon *M. thermautotrophicus* ΔH. Thermally activated graphite felt anodes were included with three different conditions (addition of Ti-NTA with L-cysteine, addition of 0.1 g·L$^{-1}$ $Na_2S$ without L-cysteine, and control with L-cysteine). Ti-NTA was supplied at a ratio of 1:1000 at the beginning and every sampling point. Each condition included iron additions of 0.1 g·L$^{-1}$. Carbon cloth was used as the cathode: **(**Y-axis 1) cathode-based geometric $CH_4$ production rates; and **(**Y-axis 2**)** Coulombic efficiencies; (**B**) Sulfide concentrations during the addition of 0.1 g·L$^{-1}$ $Na_2S$ batch experiment throughout time with the thermophilic archaeon *M. thermautotrophicus* ΔH; and (**C**) ORP measurement for all three conditions throughout time with the thermophilic archaeon *M. thermautotrophicus* ΔH. The error bars show the standard deviation for the average data from triplicate systems

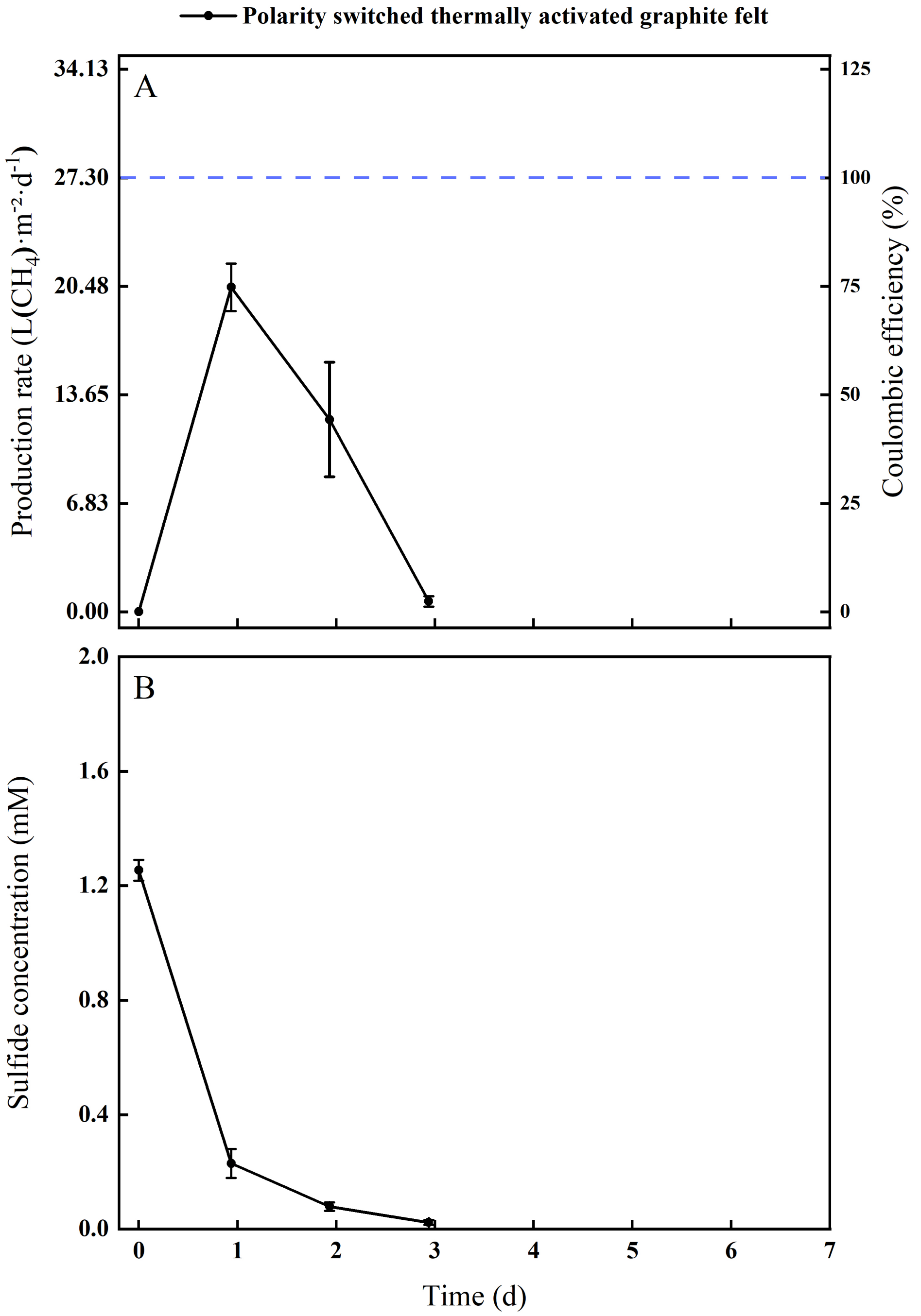


**Figure S6:** BES experiment with polarity switch: (**A**) $CH_4$ production rate corrected to the cathode surface area and Coulombic efficiency (at I = 1 $A \cdot cm^{-2}$) during batch experiments throughout time with the thermophilic archaeon *M. thermautotrophicus* ΔH when the polarity of the thermally activated graphite felt anodes was switched every 4h. 0.1 $g \cdot L^{-1}$ $Na_2S$ was supplied to the medium instead of L-cysteine; and (**B**) Sulfide concentrations during batch experiments throughout time. The error bars show the standard deviation for the average data from triplicate systems.

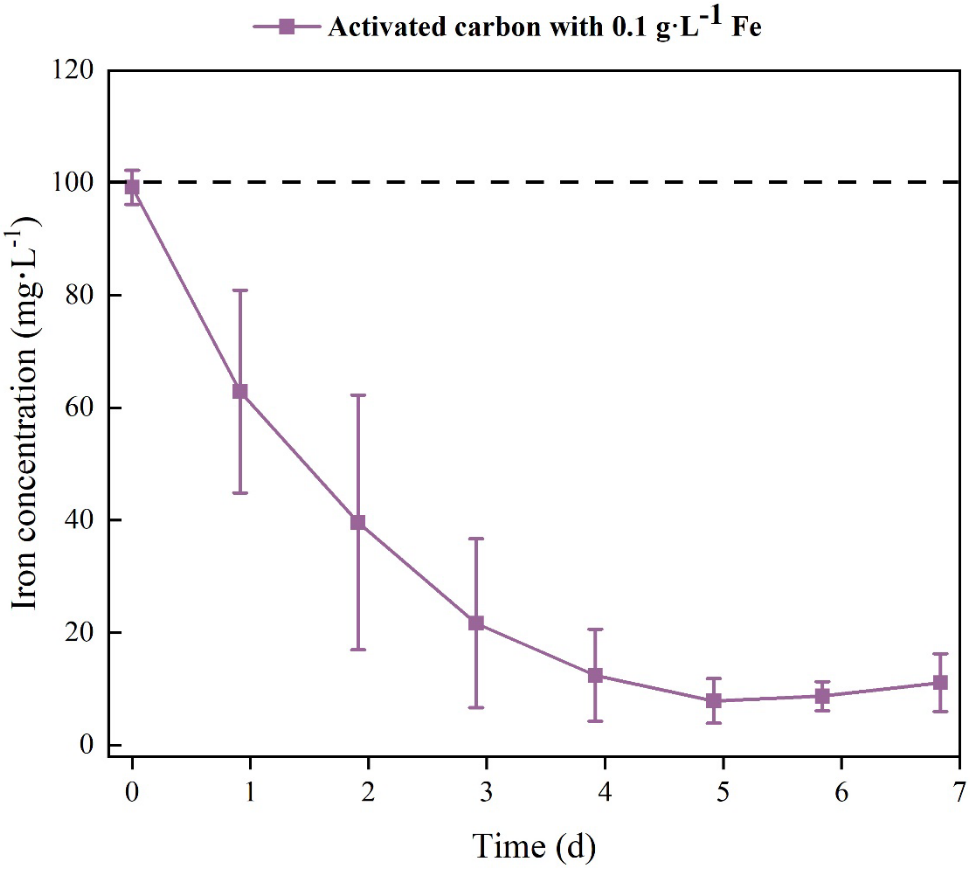


**Figure S7:** iron concentration during batch experiments of activated carbon-based anode with added iron throughout time with the thermophilic archaeon *M. thermautotrophicus* ΔH.

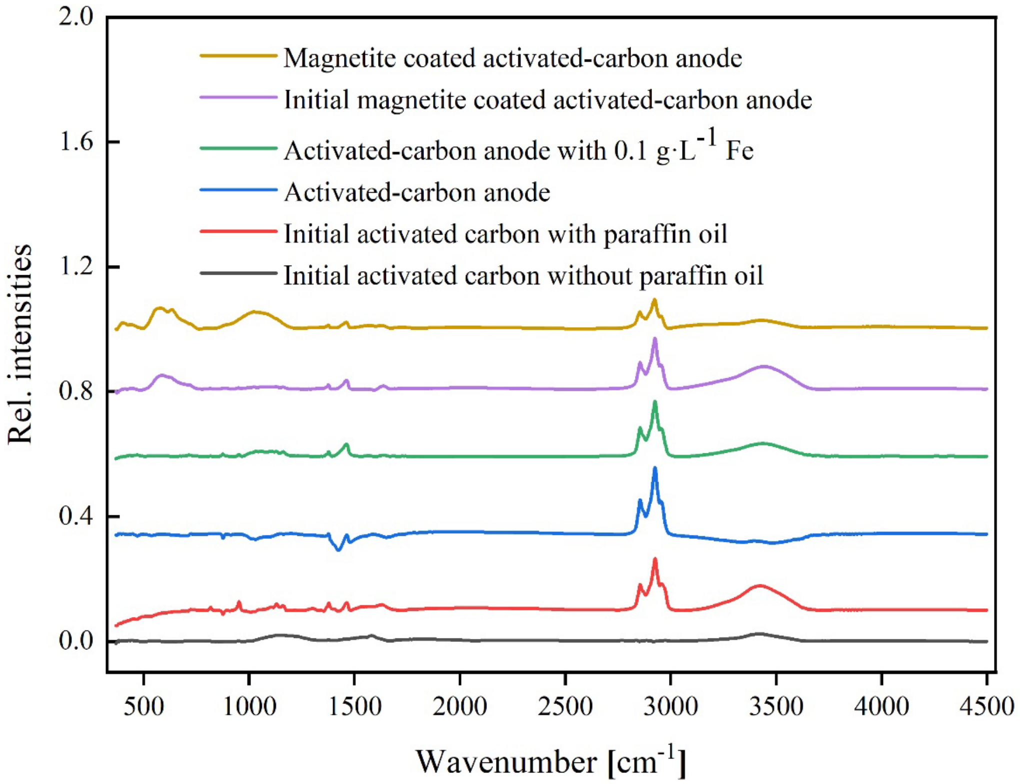


**Figure S8:** FTIR measurements of initial activated carbon with and without paraffin oil, activated-carbon anode, activated-carbon anode with 0.1 g·L$^{-1}$ Fe, initial magnetite-coated activated-carbon anode, and magnetite-coated activated-carbon anode. Samples were taken from dried anodes after 5 days of oxidation.

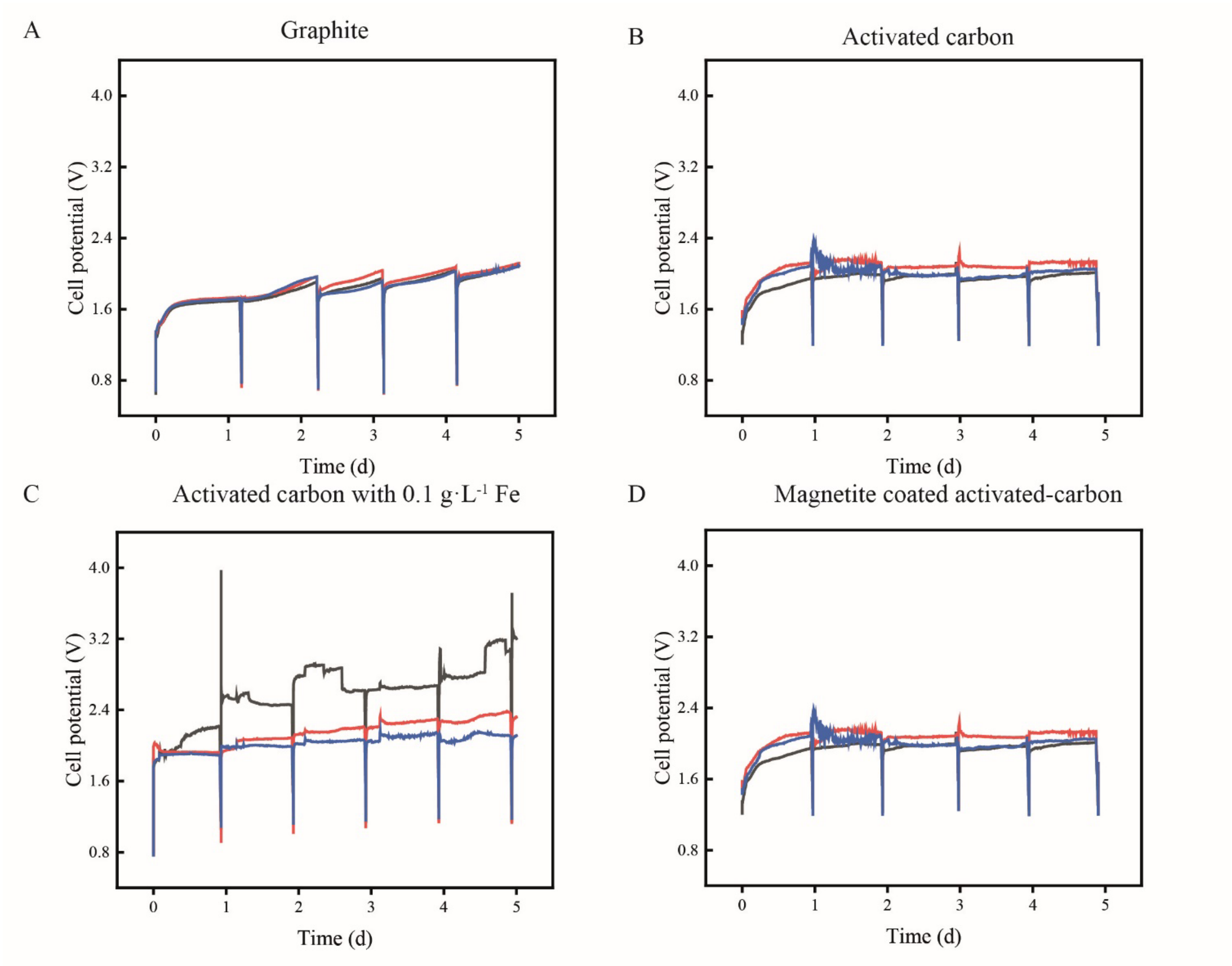


**Figure S9:** Cell potentials of the BES experiment performed in triplicates using different carbon based anodes: **(A)** graphite; **(B)** activated-carbon; **(C)** activated-carbon with 0.1 g·$L^{-1}$ Fe; and **(D)** magnetite-coated activated-carbon.

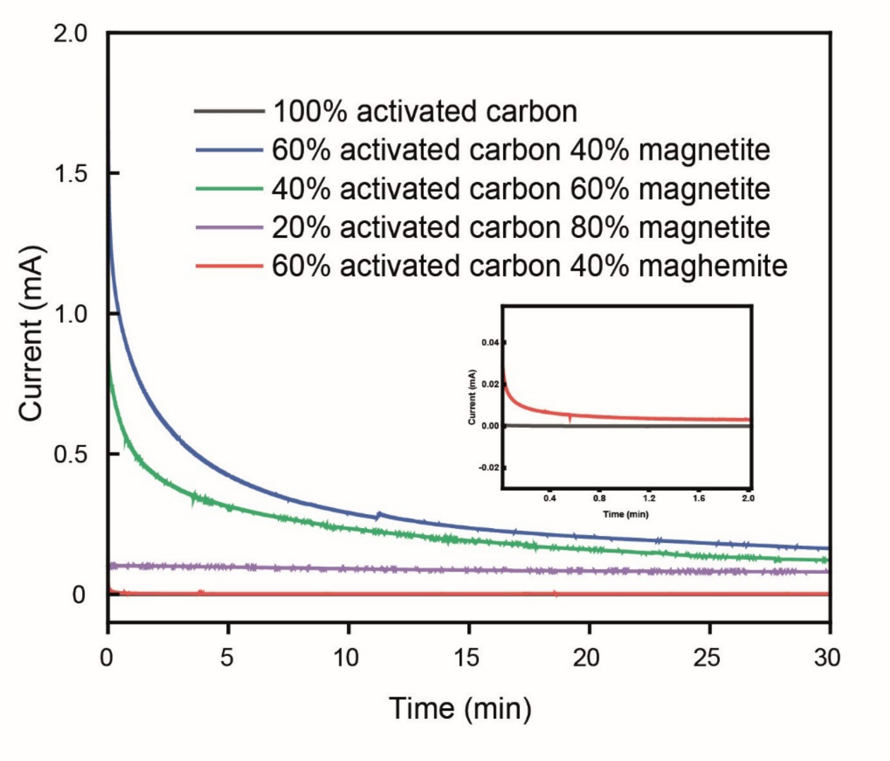


**Figure S10:** Chronoamperometic current traces of 100% activated carbon, 60% activated carbon & 40% magnetite, 40 % activated carbon & 60% magnetite, 20% activated carbon & 80% magnetite, and 60% activated carbon & 40% maghemite at +0.6 V in 66 mM phosphate buffer.

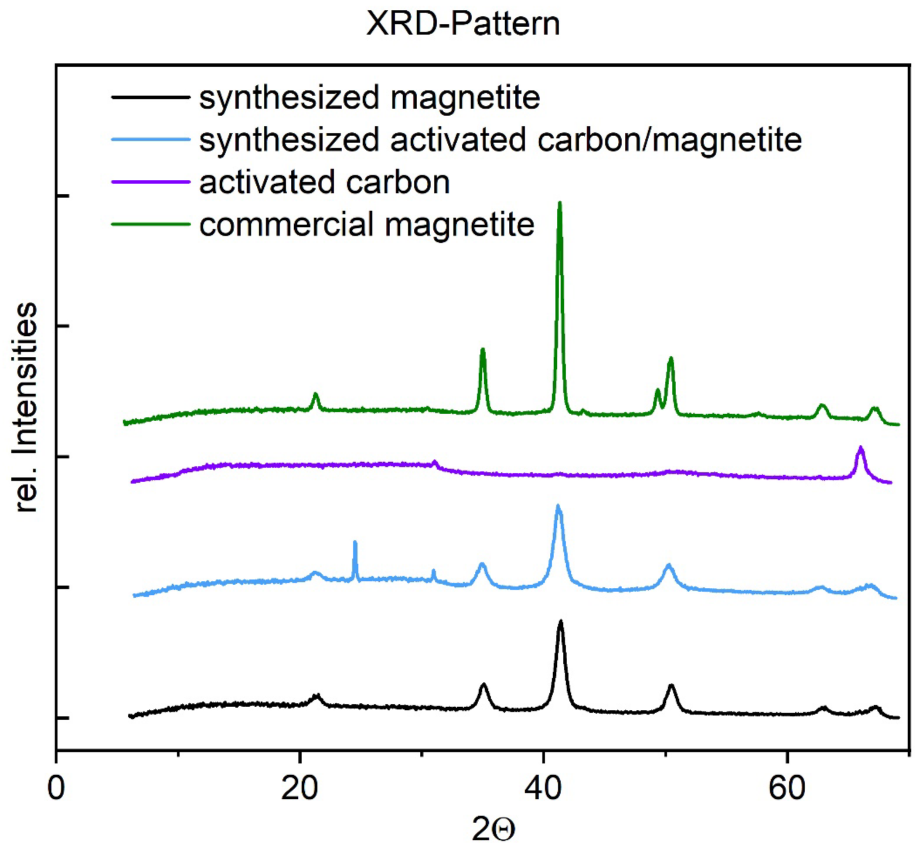


**Figure S11:** XRD pattern of synthesized magnetite, synthesized magnetite coated on activated carbon, activated carbon, and commercial magnetite.

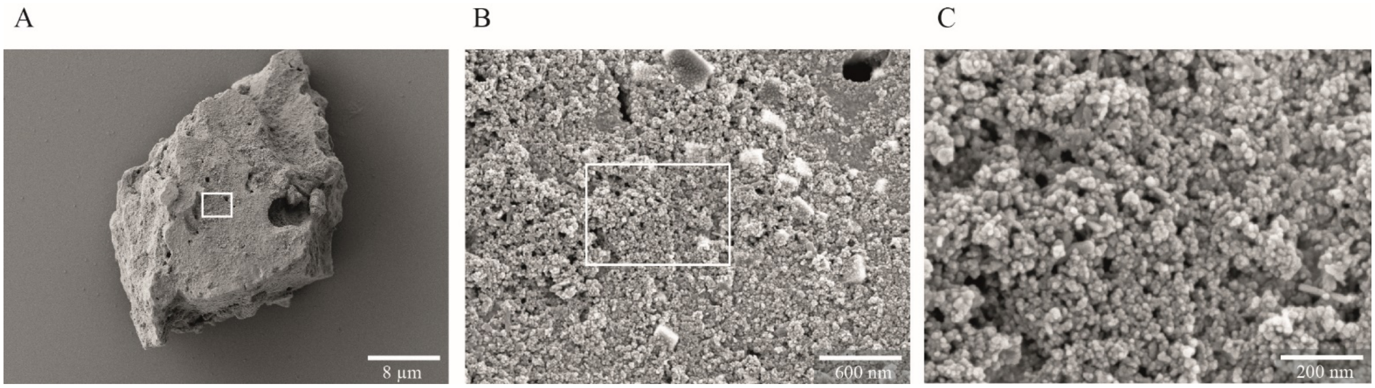


**Figure S12:** Scanning electron micrographs of magnetite-coated activated carbon: **(A)** Particle of magnetite-coated activated carbon with a 2300x magnification; **(B)** close-up from the rectangular-shaped area of (A) with a magnification of 35000x; and **(C)** close-up from the rectangular-shaped area of (B) with a magnification of 95000x.

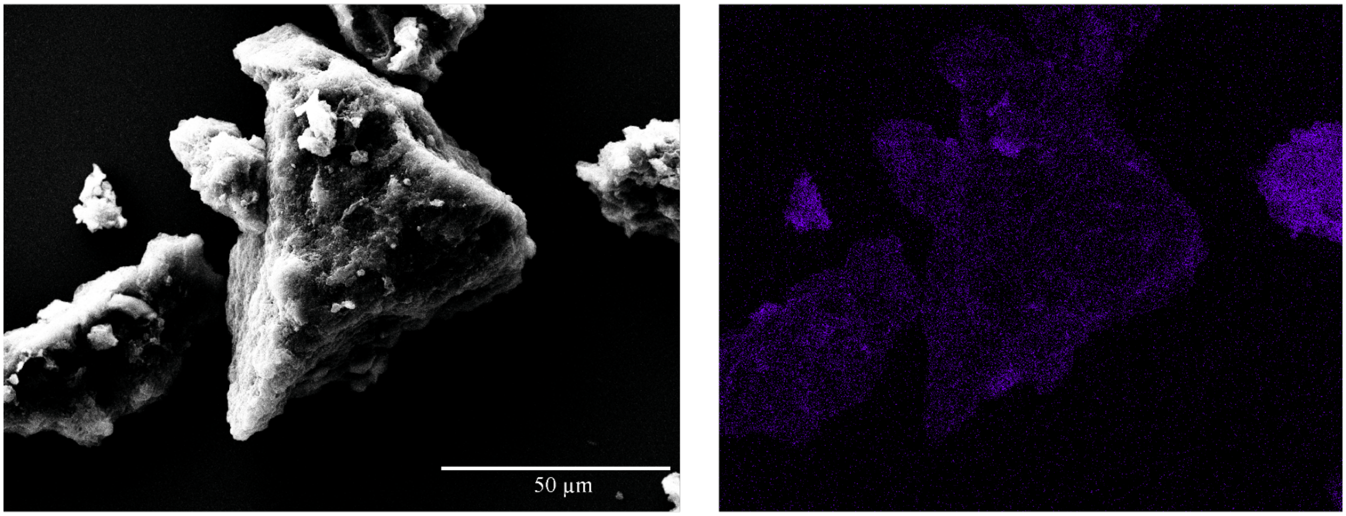


**Figure S13:** Scanning electron micrograph and its EDX Fe Kα elemental mapping (in violet).

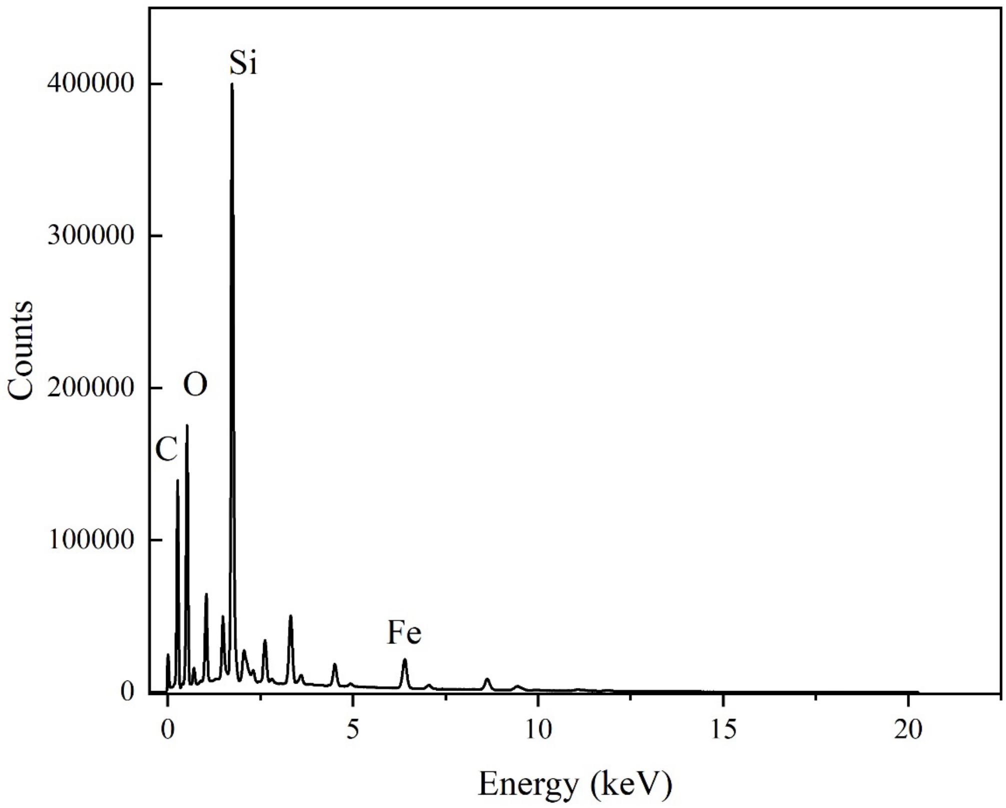


**Figure S14:** EDX Fe Kα elemental mapping spectrum from **Fig. S13**.

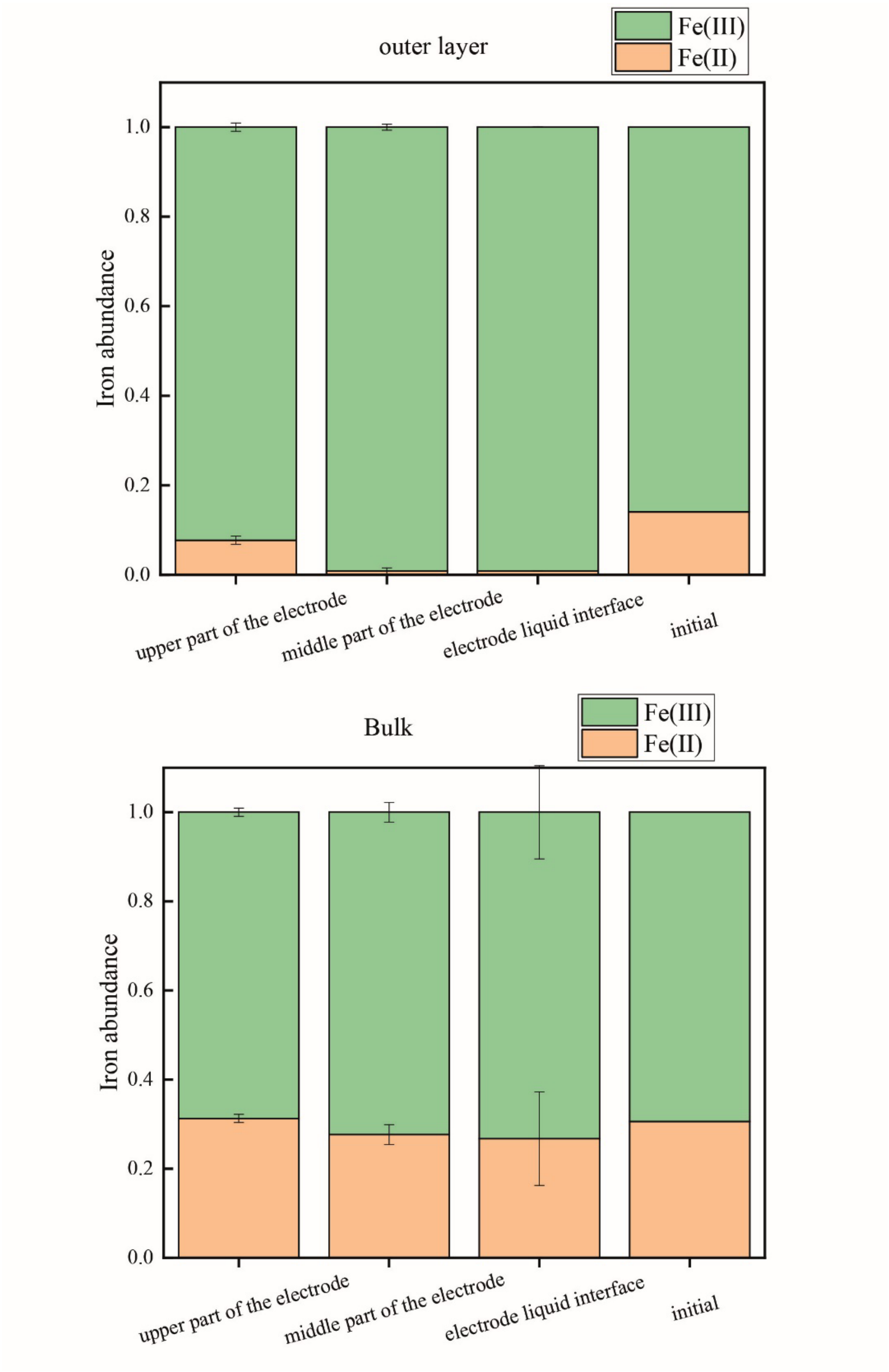


**Figure S15:** Oxidation state of magnetite. Measured Fe(II) and Fe(III) abundance from the outer layer and bulk from the upper part, middle part, and the electrode liquid interface.
